# Single-cell profiling of complex plant responses to *Pseudomonas syringae* infection

**DOI:** 10.1101/2022.10.07.511353

**Authors:** Jie Zhu, Signe Lolle, Andrea Tang, Bella Guel, Brian Kvikto, Benjamin Cole, Gitta Coaker

## Abstract

Plant response to pathogen infection varies within a leaf, yet this heterogeneity is not well resolved. We exposed *Arabidopsis* to *Pseudomonas syringae* or mock treatment and profiled >11,000 individual cells using single-cell RNA sequencing. Integrative analysis of cell populations from both treatments identified distinct pathogen responsive cell clusters exhibiting transcriptional responses ranging from immunity to susceptibility. Pseudotime analyses through pathogen infection revealed a continuum of disease progression from an immune to susceptible state. Confocal imaging of promoter reporter lines for transcripts enriched in immune cell clusters expressed surrounding substomatal cavities colonized or in close proximity to bacterial colonies, suggesting cells within immune clusters represent sites of early pathogen invasion. Susceptibility clusters exhibited more general localization and were highly induced at later stages of infection. Overall, our work uncovers cellular heterogeneity within an infected leaf and provides unique insight into plant differential response to infection at a single-cell level.

## INTRODUCTION

Plants can be infected by diverse pathogens capable of colonizing roots, vascular tissues, and foliar (leaf) tissue. Many plant diseases exhibit variable symptoms. For example, inoculation of bacteria or fungal spores using spray or infiltration results in unequal symptom development and a relatively small proportion of pathogens successfully invade their hosts (Berruyer et al., 2006; Pétriacq et al., 2016; Xin et al., 2016). Moreover, different stages of pathogen infection are often observed within a leaf (Fantozzi et al., 2021; Haueisen et al., 2019). Heterogeneity in pathogen distribution and likely, the plant response, impacts symptom development (Matsumoto et al., 2022). Strains of the bacterial pathogen, *Pseudomonas syringae*, have a broad host range and can infect many economically important plant species, causing a variety of foliar symptoms (Katagiri et al., 2002; Xin et al., 2018).

Mechanisms regulating pathogen distribution and colonization on plants can be a combination of physical, metabolic and immune barriers. Physical barriers, such as trichomes, the waxy cuticle, plant cell walls and closed stomatal pores can regulate the penetration of pathogens into the plant interior (Bigeard et al., 2015; Wang et al., 2019). Plants can also recognize pathogen molecular features, induced damage and effectors using either surface-localized or intracellular immune receptors (Yuan et al., 2021). Surface-localized pattern recognition receptors (PRRs) detect microbe-associated molecular patterns (MAMPs) or damage (DAMPs) resulting in PRR-triggered immunity (PTI). Immune recognition leads to a series of downstream defense responses including calcium influx, the production of reactive oxygen species, defense hormone production and global transcriptional reprogramming (Lolle et al., 2020; Zhou and Zhang, 2020). PTI can be generally induced against diverse pathogens because of the conserved nature of MAMPs (e.g., bacterial flagellin and elongation factor Tu, fungal chitin) (Zhou and Zhang, 2020). However, virulent pathogens secrete metabolites and proteinaceous effectors that dampen immunity and establish suitable environments for growth (Toruño et al., 2016). Thus, plant-pathogen interactions are a highly dynamic process, resulting in heterogeneous cellular responses.

Past studies investigating plant-pathogen interactions mainly depend on assays from bulk tissue (i.e. whole leaf or roots). While genome-wide transcriptional profiling has advanced our understanding of immune responses, they average cellular responses across the entire tissues (Liu et al., 2022; Zipfel et al., 2006). Single-cell RNA sequencing (scRNA-seq) technologies enable massively parallel transcriptional profiling of thousands of cells (Kolodziejczyk et al., 2015; Libault et al., 2017; Macosko et al., 2015; Tang et al., 2009). scRNA-seq interrogates populations at the single-cell level and on a genome-wide scale to profile transcriptomes from different cell types and cell states (Libault et al., 2017; Seyfferth et al., 2021). The application of scRNA-seq in plants has provided new insight into cell identity, function and development in different tissues (Denyer et al., 2019; Kim et al., 2021; Lopez-Anido et al., 2021; Procko et al., 2022; Shahan et al., 2022; Zhang et al., 2019). With respect to pathogen infection, it remains unclear how large populations of plant cells within a tissue respond and how pathogen proximity influences cellular responses at high resolution. In this study, we combined scRNA-seq and live-cell imaging of fluorescent reporters to investigate plant cellular responses to pathogen infection. We established a transcriptome atlas of *Arabidopsis* leaf tissue infected with virulent *P. syringae*. The atlas enabled the identification of pathogen-responsive cell clusters at immune, transition and susceptible states. Pseudotime trajectory revealed a continuum of disease progression from an immune to susceptible state. We validated this trajectory using fluorescent transcriptional reporter lines expressing either immune or susceptible cell cluster markers identified by scRNA-seq. Finally, we identified diverse spatial and temporal patterns of immune and susceptible marker genes which can be influenced by pathogen proximity.

## RESULTS

### Single-cell RNA-seq profiling of *Arabidopsis* leaf tissue infected with *P. syringae*

To investigate plant responses to pathogen infection at high resolution, we first analyzed bacterial distribution within a leaf. We compared the bacterial distribution between wild-type *P. syringae* pv. tomato DC3000 3xmCherry with the DC3000 *ΔhopQ1* 3xmCherry at 0, 4, 20, and 24h post-inoculation (hpi, Figure S1). Both bacterial strains behaved identically. The *hopQ1* effector deletion strain frequently used as a tool to investigate *P. syringae,* is fully virulent on *Arabidopsis* and also infects *Nicotiana benthamiana* (Wei et al., 2007). Therefore, we used virulent *P. syringae* pv. tomato DC3000 *ΔhopQ1* labeled with 3xmCherry (*Pst* DC3000, hereafter) for our experiments. *Pst* DC3000 was inoculated on four-week-old *Arabidopsis thaliana*. We observed patchy distribution of *Pst* DC3000 as well as differences in colony number and area within a leaf at 24hpi, suggesting bacterial colonization is spatially variable (Figure 1A-C). At this infection stage, *Arabidopsis* leaves do not exhibit visible symptoms, but bacteria multiply aggressively (Katagiri et al., 2002; Whalen et al., 1991).

**Figure 1.**
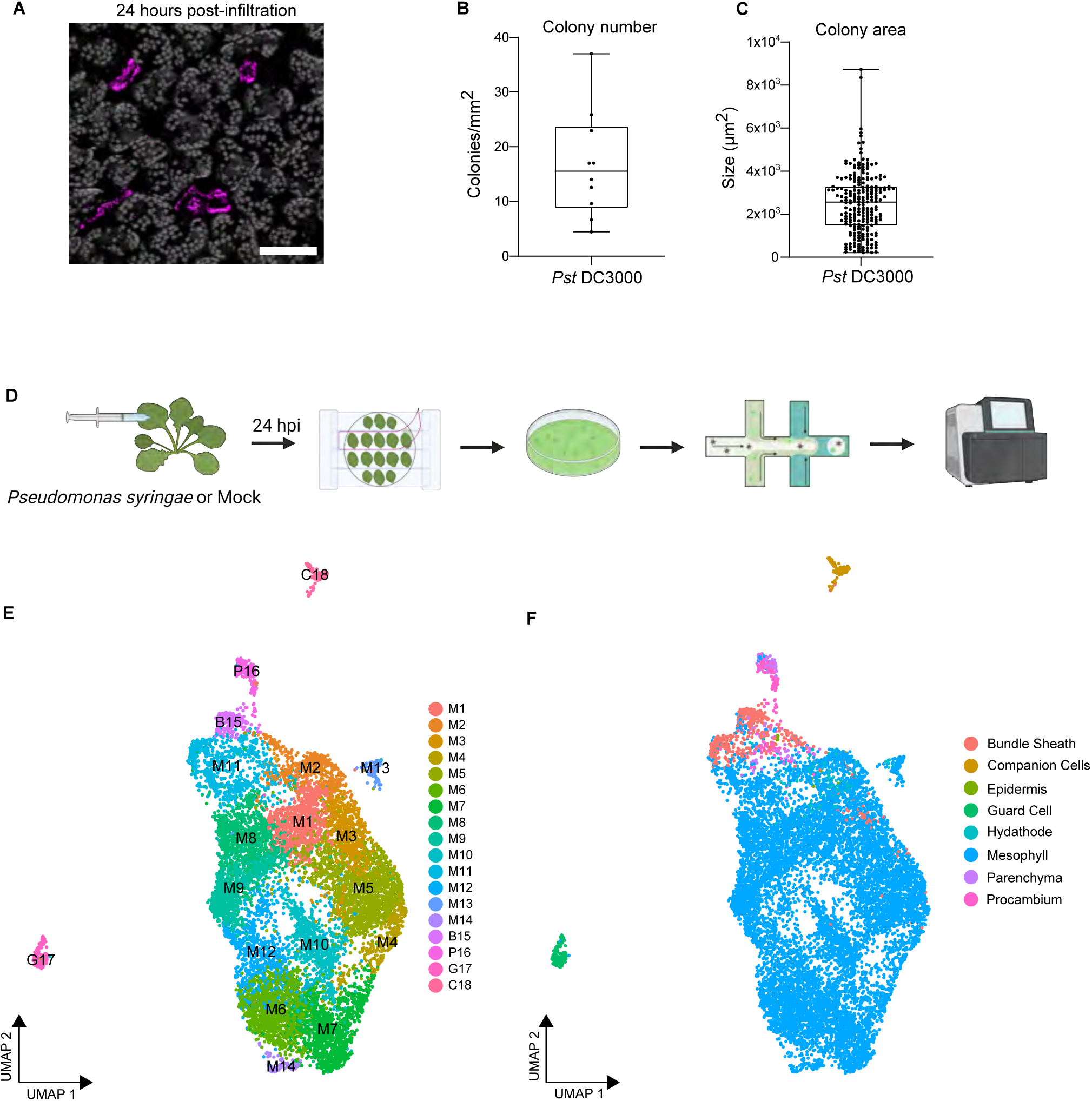
Single-cell RNA-seq profiling of *Arabidopsis* infected with Pseudomonas syringae. (A) Confocal micrograph of an *Arabidopsis* leaf 24 hours post-infiltration (hpi) with mCherry-tagged *Pst* DC3000 Δ*hopQ1* (*Pst* DC3000). The image is a maximum projection from 21 confocal Z-stacks. Chlorophyll autofluorescence is shown in grey. Scale bar = 100 µm. (B and C) *Pst* DC3000 colony number and area in infiltrated leaf tissue shown in (A). Boxplot shows median with minimum and maximum values indicated (n = 10 images from 4 plants). (D) Overview of the scRNA-seq experiment. Four-week-old *Arabidopsis* Col-0 was infiltrated with *Pst* DC3000 or 10mM MgCl_2_. Twenty-four hpi, protoplasts were prepared using the Tape-*Arabidopsis* Sandwich method. Cells were isolated on the 10x genomics Chromium chip and sequenced using the Illumina NovaSeq6000 platform. (E and F) Single Cell Uniform Manifold Approximation and Projection (UMAP) plots from both *Pst* DC3000 and mock-treated samples, colored according to cluster identities (E) and cell types (F). M: mesophyll, B: bundle sheath, P: parenchyma and procambium, G: guard cells, C: companion cells. See also Figure S1-2 and Table S1, S6.

*Pst* DC3000 colonizes the intercellular space between mesophyll cells, manipulating them to provide more favorable conditions for microbial growth. To characterize the dynamics of the interaction between *Pst* DC3000 and *Arabidopsis* mesophyll tissue, we enriched for mesophyll cells using the Tape-*Arabidopsis* Sandwich method (Figure 1D) (Wu et al., 2009). Single-cell transcriptomes were then profiled from *Pst* DC3000- and mock-treated samples 24hpi using the 10X Genomics scRNA-seq platform (Figure 1D). We recovered 11,895 single-cell transcriptomes with a median number of 3,521 genes and 17,017 unique transcripts, representing over 80% of protein-coding genes in the *Arabidopsis* genome (Figure S2A-B). Complementing our single-cell datasets, we also performed bulk RNA-seq for protoplasts and infiltrated leaves to identify genes modulated in response to pathogen infection (n = 890; adjusted p-value < 0.01, log fold-change > 2), as well as genes possibly affected by protoplast generation (n = 7548; adjusted p-value < 0.01, log fold-change > 0.5, Table S1). There was a strong correlation between merged single-cell and bulk protoplasts samples (Spearman’s Rho=0.786 and 0.848 for mock- and bacteria-treated samples, respectively, Figure S2C), indicating that protoplasting did not severely impact most genes’ expression. We excluded protoplast-inducible genes from further analysis of our single-cell dataset.

Using graph-based unsupervised clustering, we identified 18 major cell clusters and visualized them on a uniform manifold approximation and projection (UMAP) plot (Figure 1E). Each cluster contained cells from both *Pst* DC3000- and mock-treated leaves (Figure 1E and Figure S2D). To assign cell types, we utilized a recently published single-cell transcriptomics survey of *Arabidopsis* leaf tissue (Kim et al., 2021), as well as expression of well-known cell-type markers. The predominant predicted identity of cells within each cluster was then used to assign a cell type to the whole cluster. We also integrated our single-cell datasets with five previously published *Arabidopsis* leaf scRNA-seq datasets (Kim et al., 2021; Liu et al., 2020; Lopez-Anido et al., 2021; Procko et al., 2022; Zhang et al., 2021). Analysis of the integrated dataset suggests that the vast majority of cells profiled in this study are similar in cell type as those profiled by others, with the exception of an increased density of *Pst* DC3000-treated cells within the larger mesophyll cell cluster (Figure S2F-N). The 18 cell clusters contain eight cell types, but exhibit predominant mesophyll identity (∼93.7% of all cells, clusters M1 - M14, Figure 1E-F and Figure S2D).

### scRNA-seq reveals cell clusters ranging from immunity to susceptibility within a leaf

Integrative analysis of cells from *Pst* DC3000- and mock-treatment revealed a large subpopulation of cells from infected leaves comprising clusters M1 to M5 (Figure S2D and Figure S3A-B). More than 70% of cells in clusters M1-M5 were exposed to pathogen treatment, representing 34.8% of all cells. Further examination revealed that expression of genes induced by bacterial infection in these five clusters were generally higher than other clusters (Figure S3C). We refer to these five clusters, M1-M5, as pathogen-responsive clusters. Some of the cells in pathogen-responsive clusters were from the mock-treated sample. The plants used for this study were grown on soil (not in axenic conditions) and we hypothesize soil-borne or environmental bacteria that might elicit a defense response from the plant as previously described (Saarenpää et al., 2022). We examined whether impacts from protoplasting could have resulted in segmentation of this cell population. Here, we derived a protoplast signature score representing scaled expression across the set of genes found to be induced by protoplasting (Figure S2E). While some cell clusters exhibited relatively high protoplast-related expression, this could not completely explain the separation of cells from treated and untreated populations (Figure S2E). Taken together, our results indicate major differential shifts in the plant cell response after pathogen exposure.

To investigate the transcriptional reprogramming occurring in each pathogen-responsive cluster, we carried out Gene Ontology (GO) analyses. Clusters M1 and M2 exhibited enrichment of GO terms related to defense response to bacterium, immune response, and response to salicylic acid (SA) (Figure S3D). Clusters M4 and M5 were enriched in terms related to response to jasmonic acid and water transport (Figure S3D). These results suggest opposite transcriptional responses in clusters M1-M2 versus M4-5. To confirm this result, we calculated immune and susceptibility response scores based on gene expression modules for sets of genes known to be involved in immunity or disease and were differentially expressed in our bulk RNA-seq analysis (Figure 2A, Table S2). Consistent with the GO analyses, clusters M1 and M2 displayed a higher immune response score, while M4 and M5 displayed a higher susceptibility response score (Figure 2A). Cluster M3 did not display a strong average response score (Figure 2A).

**Figure 2.**
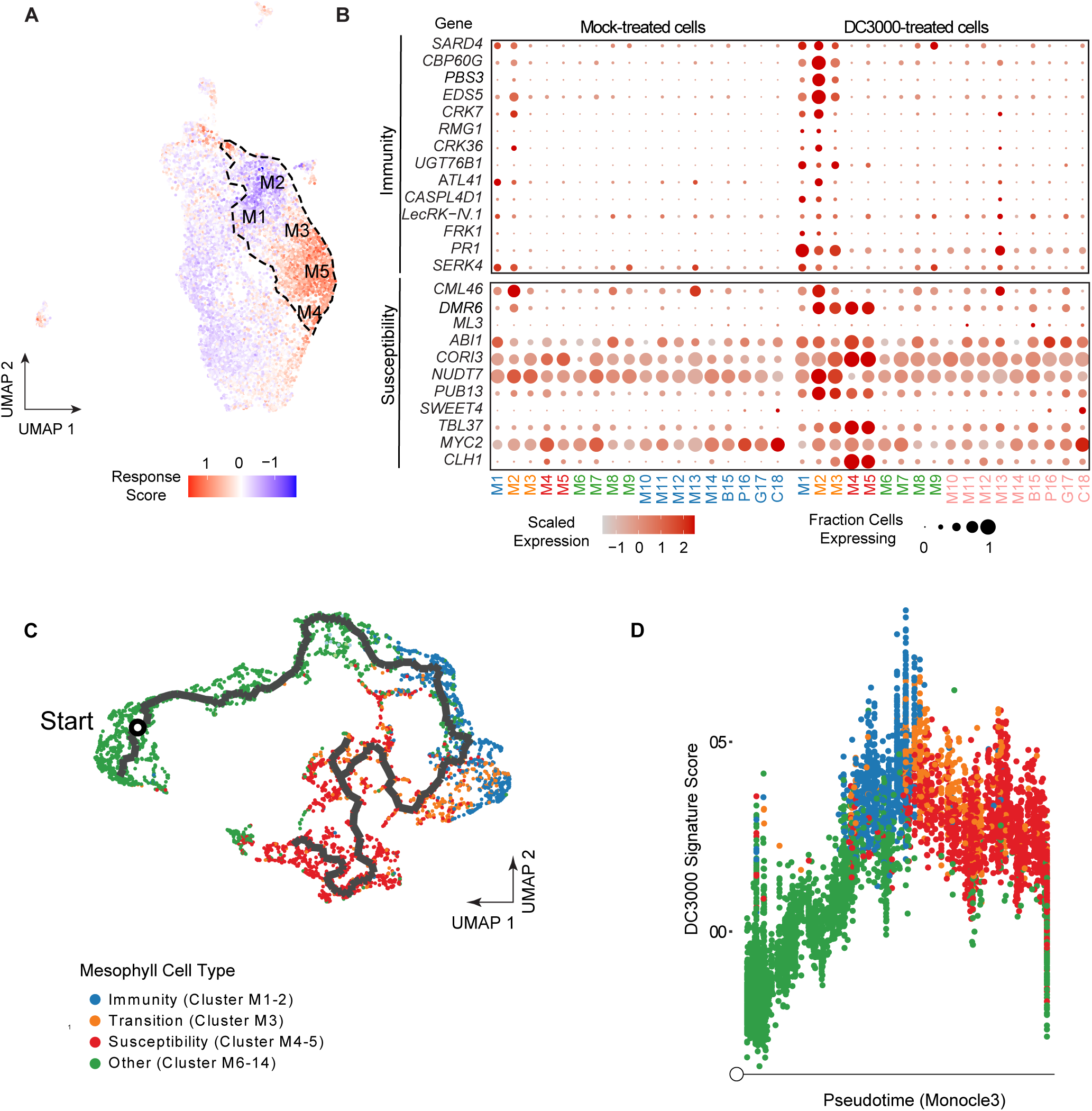
scRNA-seq reveals a continuum of disease progression from immune to susceptible responses within leaf tissue. (A) UMAP plot visualizing the magnitude of response to pathogen infection. Dashed line outlines the pathogen-responsive clusters (M1-5). Immune and susceptibility response scores were calculated as gene expression modules (Methods) based on the cell-specific expression of sets of genes known to be involved in immunity or susceptibility that were differentially expressed in our bulk RNA-seq analysis. Blue (negative values) indicates more immune-like, red (positive values) indicates more susceptible-like. (B) Dot plot of the relative expression and percent of cells expressing known plant immunity or susceptibility genes across different mesophyll cell populations in the integrated scRNA-seq data at 24hpi with *Pst* DC3000. (C) Pseudotime trajectory through mesophyll cells shows directed transition from immunity (M1-2) to susceptibility (M4-5). Mesophyll cells in clusters M1-14 were re-embedded in low-dimensional space, then subjected to trajectory inference using the Monocle3 package (Methods). An initial cell was chosen as having the lowest DC3000-induced expression signature (D). Cells colored by their cluster membership, with green cells belonging to non-responsive mesophyll cells (Other), blue cells belonging to immune clusters, orange cells corresponding to the transition cluster, and red cells belonging to susceptibility clusters. (D) DC3000-induction signature throughout pseudotime. Pseudotime values computed from the trajectory shown in (C). A DC3000-signature score was defined as the module score (Methods) for genes repressed by DC3000 from our bulk RNA-seq analysis, subtracted from the module scores for those genes that were induced. Cells are colored as in (C), based on their cluster membership. See also Figure S2-5 and Table S1-4.

Next, we analyzed the expression of known genes involved in immunity and susceptibility to *Pst* DC3000 (Figure 2B). Expression of known plant immune-related genes *CALMODULIN BINDING PROTEIN 60g* (*CBP60g*), *ENHANCED DISEASE SUSCEPTIBILITY 5* (*EDS5*), *FLG22-INDUCED RECEPTOR-LIKE KINASE 1* (*FRK1*) and *PATHOGENESIS-RELATED 1* (*PR1*) was induced in clusters M1-M2 compared to non-responsive mesophyll cell clusters (M6-M14). In contrast, clusters M4-5 displayed induced expression of genes in susceptibility including those responded to jasmonic acid (JA) and abscisic acid (ABA)(e.g., *CHLOROPHYLLASE 1/CORONATINE INDUCED 1* About 30% of cells in pathogen-responsive clusters were from mock-treated sample. The plants used for this study were grown on soil (not in axenic conditions) and we hypothesize soil-borne or environmental bacteria that might elicit a defense response from the plant as previously described (Saarenpää et al., 2022). *CORI1*), *CORONATINE INDUCED 3* (*CORI3*) and *ABA INSENSITIVE 1* (*ABI1*)). *Pst* DC3000 induces JA signaling through production of coronatine, but protoplasting can also induce JA expression (Cheong et al., 2002; Reymond et al., 2000; Zheng et al., 2012). While pathogen responsive clusters had low protoplast signature scores, other clusters with JA GO term induction had higher protoplast induced scores, which may be the result of incomplete gene removal (Figure S2E, Figure S3D). Specific transcripts related to both immunity and susceptibility were induced in cluster M3, but at lower magnitude compared to clusters M1-M2 and M4-M5 (Figure 2A and Figure S4). Cluster M2 exhibited the strongest activation of immune-related genes and related biological processes (Figure 2A-B and Figure S3D). These data indicate that M1-M2 represent immune activated clusters, M3 is a transition cluster, and M4-M5 represent susceptibility clusters (Figure 2A-B). *Pst* DC3000 is able to cause disease on *Arabidopsis* and consistent with a compatible interaction, M5 represents the largest cluster (14.2% of 11, 895 cells, Figure S2D).

Within a tissue, there are multiple points of infection representing different stages of disease development. Pseudotime analyses, which aim to order cells relative to a temporal, developmental or treatment axis, have been used to model the trajectory of a biological process, with each cell signifying a singular time point along a continuum (Trapnell et al., 2014). In order to predict the trajectory of pathogen-responsive cellular clusters, we performed pseudotime analysis using Monocle3 (Cao et al., 2019; Qiu et al., 2017b, 2017a; Trapnell et al., 2014). To circumvent influence of different cell types, we used *Pst* DC3000-treated mesophyll cells from clusters M1-M14 to infer the trajectory of disease progression. The trajectory was mostly linear, progressing through non-pathogen responsive clusters (M6-14), followed by immune clusters (M1-2), the transition cluster M3 and ended in the susceptible clusters (M4-M5, Figure 2C). In order to investigate pathogen responsiveness through pseudotime, a signature score was computed to quantify the overall impact *Pst* DC3000 has on each cell utilizing module scores from genes identified as differentially expressed in our bulk RNA-seq data. When overlaid upon pseudotime, the *Pst* DC3000 signature score was markedly induced in cells undergoing immunity (clusters M1-2), then plateaued or decreased through cells experiencing features associated with disease susceptibility (clusters M3-M5), consistent with the dynamic nature of plant-pathogen interactions (Figure 2D). These results indicate in a compatible interaction, disease progresses from plant defense, into a transitional state and culminates in susceptibility.

We also sought to identify genes that have dynamic expression patterns relative to the imputed pseudotime axis. Here, we again used monocle3 to identify genes that vary significantly with pseudotime, retrieving 776 loci (Figure S5 and Table S3). We clustered these into seven groups, with two clusters having expression profiles consistent with a susceptibility response (clusters 2 and 6), while four clusters had profiles consistent with immunity or the transition between immunity and susceptibility (clusters 3-5 and 7) (Figure S5 and Table S3). Genes within clusters consistent with a susceptibility or immune response represent candidates important in disease progression (Figure S5). We also identified unique transcripts expressed in immunity, transition and susceptibility clusters (Figure S4).

### Visualization of immune and susceptible cellular markers during disease progression

We sought to experimentally validate pseudotime predictions and investigate expression of immune and susceptible markers through the course of infection. We utilized surface inoculation, which more closely mimics natural infection, on two-week-old *Arabidopsis* seedlings. This protocol also allows for inoculation of younger plants to facilitate imaging by confocal microscopy. In order to test the experimental setup, we first monitored the growth of *Pst* DC3000-mCherry (Figure 3A). Bacterial titers did not change in the first 4h post-inoculation (hpi), but dramatically increased from 24-72hpi, consistent with previous experiments on seedlings and soil-grown plants (Figure 3A left) (Katagiri et al., 2002 and Ishiga et al., 2011). We visualized spatiotemporal dynamics of bacterial colonization after surface inoculation. At 24hpi, spotty fluorescence signals were observed (Figure 3A middle), which was similar to syringe-infiltration (Figure 1A). Visualizing multiple focal planes of bacterial fluorescence signals demonstrated that bacteria colonized intercellular space between mesophyll cells by 24hpi. At 48 and 72hpi, we observed enlarged and merged fluorescent spots, indicating high multiplication of bacterial populations at these stages (Figure 3A middle). Quantification of fluorescence intensity per colony showed a continuous increase over time (Figure 3A right).

**Figure 3.**
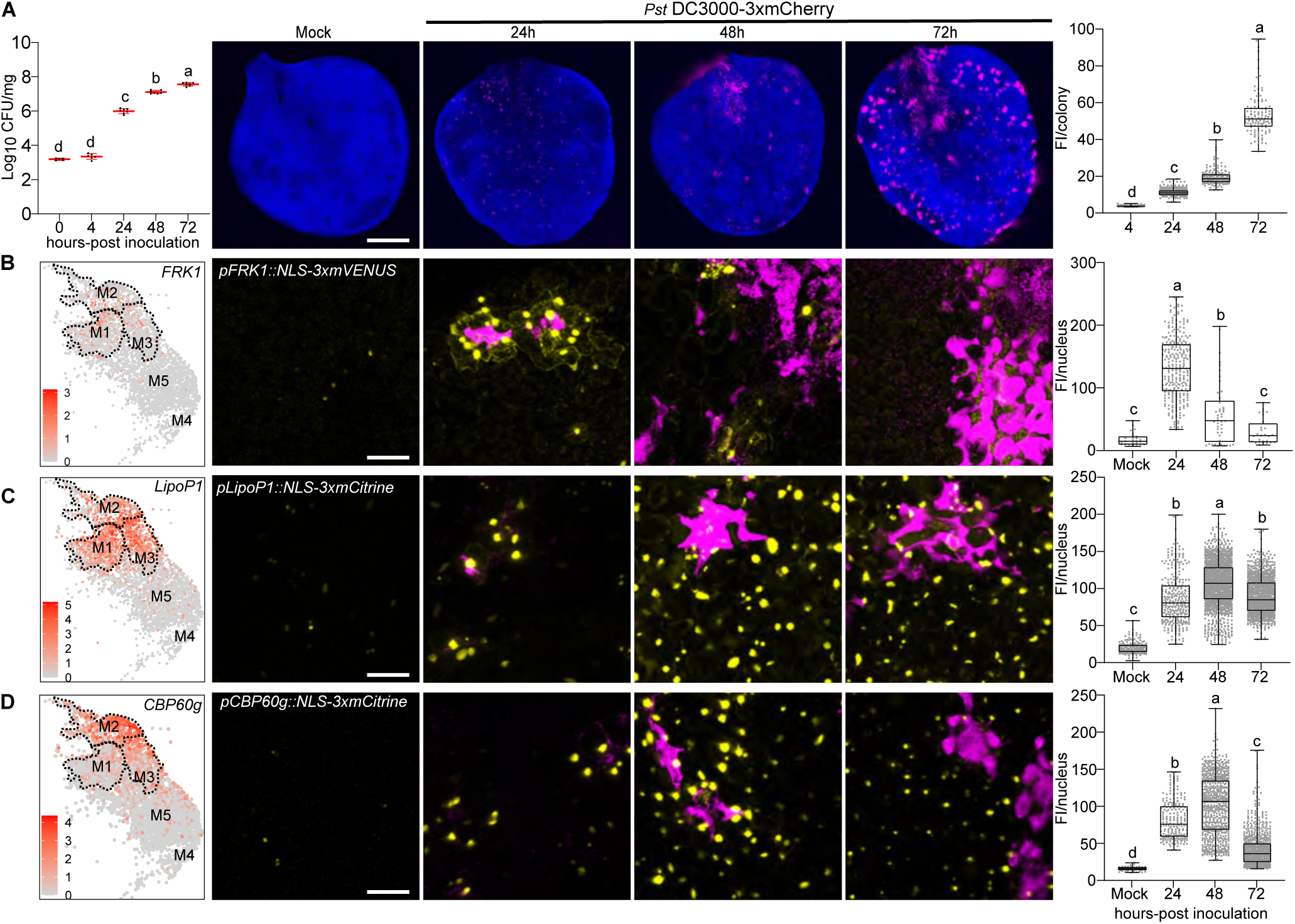
Temporal dynamics of immune marker expression during *Pseudomonas* infection. (A) Analysis of bacterial growth over three days post-flood inoculation. Two-week-old *Arabidopsis* seedlings grown on Murashige-Skoog plates were surface-inoculated with mCherry-tagged *Pst* DC3000 at concentration of 1 x 10^7^ colony forming units/ml (CFU/ml). Left: Analyses of bacterial growth over time. Middle: Confocal micrograph of representative images of mock or *Pst* DC3000-inoculated *Arabidopsis* leaves over time. Chlorophyll autofluorescence is shown in blue. Right: Bacterial populations were determined by quantifying mean fluorescence intensity (FI, mean gray values) per colony. Boxplots show median with minimum and maximum values (n = 3 images from 3 plants). Different letters indicate statistically significant differences (p < 0.0001, ANOVA with Tukey test). Scale bars: 1 mm (B, C and D) The immune markers *FRK1* (B), *LipoP1* (C) and *CBP60g* (D) are highly expressed at early infection stages but downregulated at late stages. Promoter-reporter lines for each immune marker were generated with fusion to 3xfluorophore possessing a nuclear localization signal (NLS). Left: Feature plot of immune markers in pathogen-responsive clusters. Dashed line outlines Clusters M1-M3. Middle: Representative images of immune marker expression at different infection stages. Plants were inoculated as described in (A) and mock images are taken at 24h. Pictures are maximum projections from confocal Z stacks. Right: Mean florescence intensity per nucleus was calculated and boxplot shows median with minimum and maximum values (n = 6 images from 3 plants). Different letters indicate statistically significant differences (p < 0.0001, ANOVA with Tukey test). Scale bars: 50 µm. All experiments were repeated at least two times with similar results. See also Figure S6.

We then visualized the expression patterns of cell cluster marker genes and fluorescently tagged *Pst* DC3000 during the course of infection. We selected marker genes that showed relatively high and specific expression from scRNA-seq analyses in either immune or susceptible clusters. Fluorescent transcriptional reporter lines coupled to a nuclear localization signal (NLS) enabled visualization of cell specific plant responses during pathogen infection.

Three immune marker genes were selected: *FRK1, AT3G18250* (*LIPOPROTEIN 1, LipoP1*) and *CBP60g. FRK1* is a receptor-like kinase that is strongly induced during pattern-triggered immunity and at early stages after pathogen infection (Asai et al., 2002; He et al., 2006). *LipoP1* is a putative membrane lipoprotein, whose function is unknown. The *CBP60g* transcription factor regulates biosynthesis of the plant defense hormone salicylic acid (Kim et al., 2022; Zhang et al., 2010). All three markers displayed high expression in immune clusters M1-2, variable expression in the transition cluster M3, but low expression in susceptible clusters M4-5 (Figure 3B-D left).

Using a previously characterized transcriptional reporter *pFRK1::NLS-3xmVENUS* (Zhou et al., 2020) and newly generated transgenic lines *pLipoP1/pCBP60g::NLS-3xmCitrine*, we observed low fluorescence signals in mock-inoculated *Arabidopsis* true leaves (Mock, Figure 3B-3D). A time course experiment was able to detect *FRK1* expression at earlier time points (4h and 10h), but the induction was weaker and not significant from the mock inoculation (Figure S6A). *FRK1* expression was strongly induced at 24hpi and dramatically downregulated at 48 and 72hpi (Figure 3B). *FRK1* was previously shown to be strongly induced 2h post-syringe infiltration with *Pst* DC3000 using qPCR (He et al., 2006). These results suggest that the expression of *FRK1* at 24hpi represents an early infection stage using surface inoculation. We observed the expression of *LipoP1* and *CBP60g* was highly induced at 24-48hpi (Figure 3C and 3D). All three immune markers *FRK1, CBP60g* and *LipoP1* exhibited more localized induction at 24hpi. The expression of all immune markers was significantly reduced at 72hpi when plants exhibited chlorosis and water-soaked symptoms (Figure 3B-D). A second independent transgenic line of *pLipoP1::NLS-3xmCitrine* exhibited a similar expression pattern, but had more robust expression at 72h (Figure S6C). Compared to *CBP60g* and *FRK1, LipoP1* exhibited stronger expression in the transition cluster M3, which might result in stochasticity of the late expression of this gene in different lines (Figure 3C, Figure S6C). We screened four transcriptional reporter lines of *CBP60g* (Figure S7F). Three of them (22-1, 22-18, 22-23) had similar pattern of expression and exhibited localized expression surrounding bacterial colonies at 24hpi. In contrast, one *CBP60g* line (22-4) was induced at 24h, but did not exhibit localized expression (Figure S7F). Together, these data highlight activation of immune marker genes at early infection stages and downregulation at late stages, consistent with the pseudotime trajectory.

We explored expression of three susceptibility marker genes strongly expressed in clusters M4-5: *EXPANSIN A10* (*EXPA10*), *PLASMA MEMBRANE INTRINSIC PROTEIN 1;4* (*PIP1;4*) and *IAA-leu-resistant-like5* (*ILL5*). *EXPA10* belongs to the expansin gene family whose members are able to induce cell wall loosening through a non-enzymatic function and have been implicated in plant-pathogen interactions (Cosgrove, 2015; Ding et al., 2008). *PIP1;4* is a plasma membrane localized aquaporin. Of the 13 *PIP* family members in *Arabidopsis*, 10 were significantly induced in clusters M4-M5. Aquaporins are membrane channels that facilitate the transport of water and small neutral molecules (H_2_O, H_2_O_2_, CO_2_) (Maurel et al., 2015). *ILL5* is most similar to *ILL3,* which encodes an amidohydrolase, involved in converting indole-3-acetic acid (IAA) from an amino acid conjugate to a free form to increase auxin signaling (Casanova-Sáez and Voß, 2019; Hayashi et al., 2021). All three markers displayed high expression in susceptibility clusters M4-5 (Figure 4A-C, left). *EXPA10* and *ILL5* had relatively weak expression in the transition cluster M3 and low expression in immune clusters M1-M2 (Figure 4A, 4C). In contrast, *PIP1;4* was expressed in clusters M1-M3, but at a lower level than M4-5 (Figure 4B).

**Figure 4.**
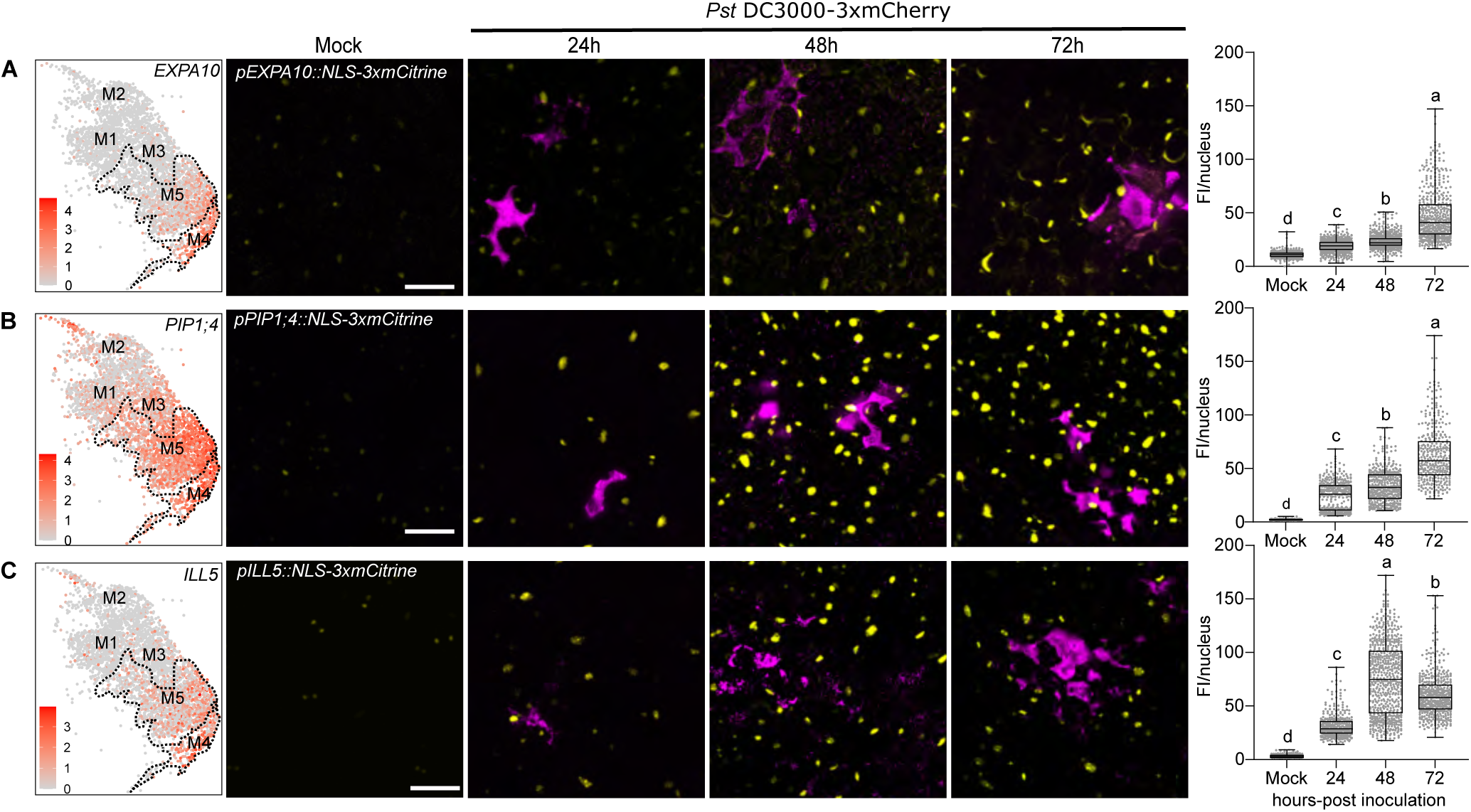
Temporal dynamics of susceptible marker expression during *Pseudomonas* infection. (A, B and C) The susceptible markers *EXPA10* (A), *PIP1;4* (B) and *ILL5* (C) are highly induced at late infection stages. Promoter-reporter lines for each immune marker were generated with fusion to 3xfluorophore possessing a nuclear localization signal (NLS). Left: Feature plot of susceptible markers in pathogen-responsive clusters. Dashed line outlines clusters M4-M5. Middle: Representative images of susceptible marker expression at different infection stages. Two-week-old *Arabidopsis* seedlings grown on Murashige-Skoog plates were surface-inoculated with mCherry-tagged *Pst* DC3000 and mock images were taken at 24h. Pictures are maximum projections from confocal Z stacks. Right: Mean florescence intensity (FI, mean gray values) per nucleus was calculated and boxplot shows median with minimum and maximum values indicated (n = 6 images from 3 plants). Different letters indicate statistically significant differences (p < 0.0001, ANOVA with Tukey test). All experiments were repeated at least two times with similar results. Scale bars: 50 µm. See also Figure S6.

We examined the expression of two independent transgenic lines for each susceptibility marker after inoculation with *Pst* DC3000 and observed similar results for both lines (*pEXPA10::NLS-3xmCitrine, pPIP1;4::NLS-3xmCitrine, pILL5::NLS-3xmCitrine,* Figure 4A-C, Figure S6D-F). The *EXPA10* and *PIP1;4* transcriptional reporter lines exhibited low levels of expression in mock-inoculated plants and increasing levels of expression after inoculation, peaking at 72hpi (Figure 4A-B, Figure S6D-E). A time course experiment was able to detect *EXPA10* induction at 10h, but at a lower level than the 24, 48 or 72hpi time points (Figure 4A, Figure S6D). The *ILL5* transcriptional reporter lines exhibited low levels of expression in mock-inoculated plants and increasing levels of expression, peaking at 48hpi (Figure 4C, Figure S6F). *ILL5* expression decreased slightly at 72hpi, but was still higher than 24hpi (Figure 4C, Figure S6F). Thus, the susceptible markers are activated after bacterial infection and strongly induced during later infection stages. These data highlight expression of susceptibility genes at later infection stages, consistent with the pseudotime trajectory.

### Immune and susceptible marker genes exhibit diverse patterns of spatial expression

Bacteria exhibit heterogeneity in colonization of a leaf after both syringe and surface inoculation (Figure 1A, Figure 3A). Our transcriptional reporter lines enabled us to probe marker gene expression with high sensitivity and at single-cell resolution. Therefore, we investigated where pathogen-responsive cells are spatially localized after surface inoculation with *Pst* DC3000. First, we examined expression of the *FRK1* immune marker gene at 24hpi when it showed highest expression (Figure 3B). *FRK1* was rarely expressed in mock samples, but was frequently observed in cells surrounding substomatal cavities colonized by bacteria (Figure 5A and Movie S1-S2). *P. syringae* uses stomatal pores to enter the leaf interior and colonize the substomatal cavity early during infection (Melotto et al., 2008). We quantified confocal micrographs to determine the spatial localization of *FRK1* expressing cells. Ninety-one percent of cells expressing *FRK1* after *Pst* DC3000 inoculation surrounded substomatal cavities (Figure 5B). Next, we quantified the proximity of *FRK1* expressing cells to bacterial colonies. Seventy-five percent of *FRK1* expressing cells were proximal (< 15 μm) to bacterial colonies (Figure 5C). These data demonstrate that *FRK1*, which is known to be one of the earliest PTI marker genes, exhibits strong and specific expression at sites of bacterial invasion.

**Figure 5.**
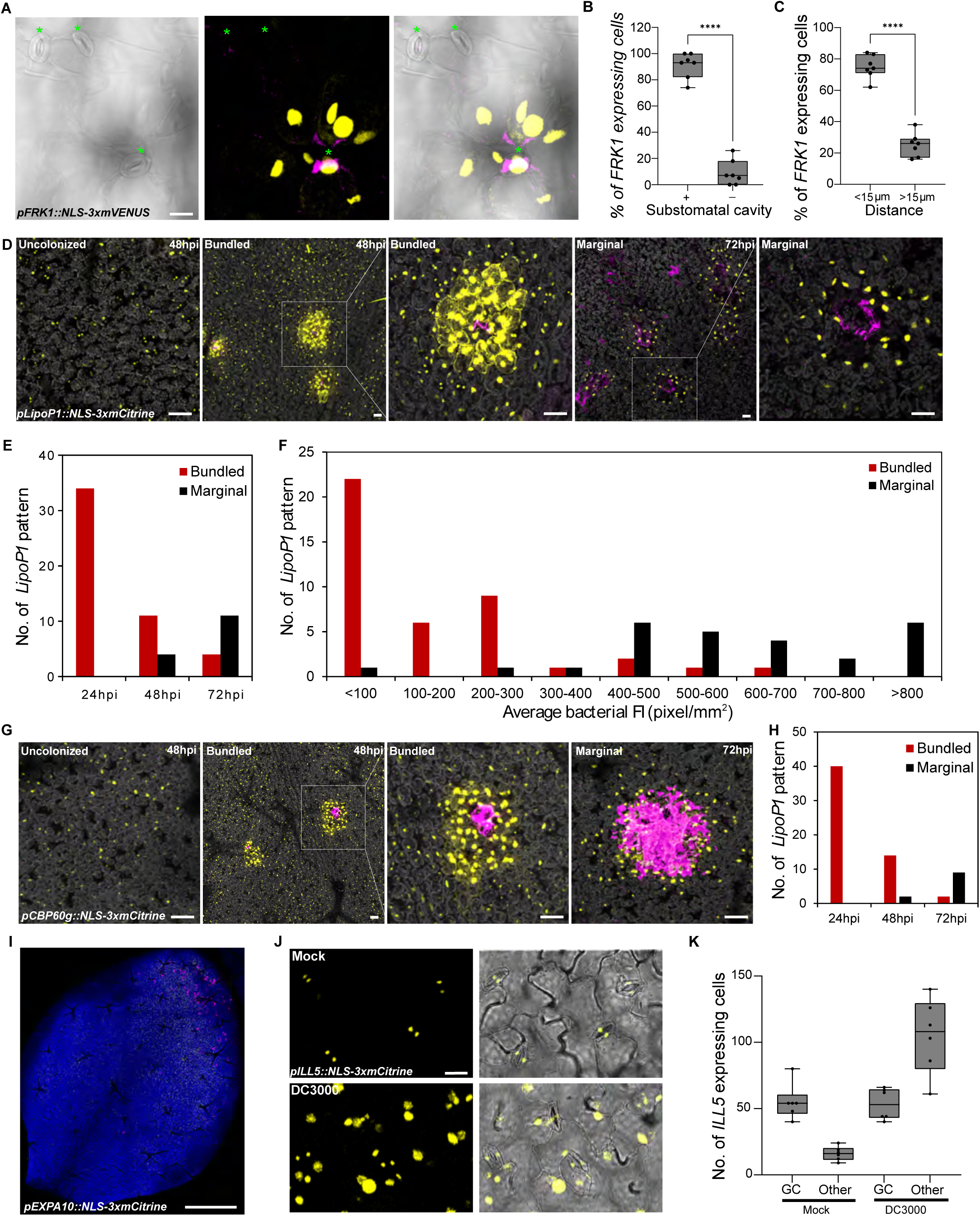
Spatial dynamics of immune and susceptible marker expression after *Pseudomonas* infection. (A-C) The immune marker *FRK1* is induced in surrounding cells of substomatal cavities colonized by *Pseudomonas syringae* DC3000. Two-week old *Arabidopsis pFRK1::NLS-3xmVENUS* seedlings were flood-inoculated with mCherry-tagged *Pst* DC3000. (A) At 24hpi, whole seedlings were fixed and cleared using ClearSee. Green asterisks indicate stomata. Left: A single image of bright field channel. Middle: Maximum projections of Z stack of mVENUS and mCherry signals. Each yellow dot indicates a single nucleus. Right: Merged image. Scale bar: 20 µm. (B) Percentage of *FRK1* expressing cells surrounding a substomatal cavity (+) or not (-) at 24hpi. Boxplot shows median with minimum and maximum values indicated (n = 7 images from 4 plants). ****p < 0.0001 analyzed by two-tailed, unpaired Student’s t-test. (C) Percentage of *FRK1* expressing cells that are proximal (<15 µm) or distal (>15 µm) to a bacterial colony 24hpi. Data was analyzed as described in (B). (D) The immune marker *LipoP1* is expressed in bundled and marginal patterns surrounding bacterial colonies. *Arabidopsis pLipoP1::NLS-3xmCitrine* seedlings were inoculated as described in (A). Representative images of *LipoP1* expression at 48 and 72h in DC3000-treated samples. Pictures are maximum projections of a Z stack of mCitrine, mCherry and chlorophyll autofluorescence signals. Chlorophyll autofluorescence is shown in gray. Scale bars: 50 µm. (E) Number of *LipoP1* expressing patterns at 24, 48 and 72h in DC3000-treated samples. At least 8 images from 3 plants were analyzed at each time point. (F) Number of *LipoP1* expressing patterns at different average florescence intensities (FI) of bacterial colonies. Twenty-four images taken at 48 and 72hpi were analyzed. (G) The immune marker *CBP60g* is expressed in bundled and marginal patterns during late infection. *Arabidopsis pCBP60g::NLS-3xmCitrine* seedlings were inoculated as described in (A). Representative images of *CBP60g* expression at 48 and 72hpi. Pictures are maximum projections of a Z stack of mCitrine, mCherry and chlorophyll autofluorescence signals. Chlorophyll autofluorescence is shown in gray. Scale bars: 50 µm (H) Number of *CBP60g* expressing patterns at 24, 48 and 72hpi. At least 7 images from 3 plants were analyzed at each time point. (I) Expression of the susceptible marker *EXPA10* in a larger area of the leaf proximal to DC3000 colonies at 24hpi. Maximum projections of Z stack of mVENUS, mCherry and chlorophyll autofluorescence signals. Scale bars: 1 mm. (J)The susceptible marker *ILL5* is expressed in guard cells before inoculation and broadly expressed after infection. *Arabidopsis pILL5::NLS-3xmCitrine* seedlings were inoculated as described in (A). Representative images of *ILL5* expression at 48h in mock- or DC3000-treated samples. Maximum projections of Z stack of mCitrine signals were combined with single image of bright filed channel. Scale bar: 20 µm. (K) Number of *ILL5* expressing cells at 24h in mock- or DC3000-treated samples. Boxplot shows median with minimum and maximum values indicated (n = 6 images from 3 plants). p < 0.0001, ANOVA with Tukey test. All experiments were repeated at least twice with similar results. See also Figure S7, Movie S1, Movie S2.

In contrast to *FRK1*, the *LipoP1* transcriptional reporter line exhibited dynamic changes in spatial expression patterns during the course of infection. *LipoP1* is expressed in guard cells, which flank stomatal pores, in the absence of pathogen infection (Figure 3C, Figure S6C). At 24hpi, 60% of *LipoP1* expressing cells were proximal to bacterial colonies (< 15 μm, Figure S7C). However, *LipoP1* exhibited variable spatial expression at 48 and 72hpi, either generally induced in all cells or highly induced in cells surrounding bacterial colonies, possibly due to unsynchronized bacterial infection within different leaves. We analyzed the expression pattern of *LipoP1* over time using bacterial fluorescence intensity as a proxy for bacterial colony size. In particular, we observed two patterns of *LipoP1* expression in cells surrounding bacterial colonies: a robust expression pattern surrounding colonies with a fluorescence intensity less than 300 pixels/mm^2^ (bundled pattern) as well as expression at the margins of larger colonies with a fluorescence intensity greater than 400 pixels/mm^2^ (marginal expression, Figure 5D-F)). Bundled and marginal *LipoP1* expression patterns were significantly higher than in uncolonized regions (Figure S7D). Plant cells at the center of marginal pattern did not exhibit chlorophyll autofluorescence, which indicates they died due to severe bacterial colonization. Collectively, these results suggest different *LipoP1* expression patterns are associated with bacterial population size.

*CPB60g* encodes a master immune transcription factor that works in parallel with SARD1 to synthesize the plant defense hormone SA (Kim et al., 2022; Sun et al., 2015). In the absence of pathogen infection, *CBP60g* is generally expressed with low levels in all cells. After *Pst* DC3000 inoculation, the *CPB60g* transcriptional reporter line exhibited similar induction patterns to *LipoP1*, including general, bundled and marginal patterns (Figure 5G-H). The general induction of *CPB60g* may be reflective of its role in SA biogenesis and regulation via NPR1 receptors, which are critical for within-leaf and systemic immune responses (Kim et al., 2022; Sun et al., 2015).

The intriguing spatial association of immune marker expression and bacterial colonization prompted us investigate if expression of susceptibility markers were spatially associated with bacterial colonization. We observed more broad induction of expression of the susceptibility marker *EXPA10* in large areas of the leaf that were proximal to regions robustly colonized by bacteria (Figure 5I), which contrasts with the more specific expression of the immune markers *FRK1,LipoP1* and *CBP60g*. This *EXPA10* pattern of induction in sections of the leaf was most striking at 24hpi before more uniform colonization of the leaf with larger bacterial colonies (Figure 3A, Figure 5I). Reporter lines for *EXPA10* and *PIP1;4* exhibited detectable expression in epidermal and mesophyll cells in the absence of pathogen infection (Figure 4, Figure S6E-F). In contrast, *ILL5* was mainly expressed in guard cells in leaves without bacterial infection, but strongly induced at 48-72hpi in epidermal, mesophyll and guard cells (Figure 5J-K). Collectively, the transcriptional reporter lines representing immune and susceptible markers reveal distinct patterns of spatial and temporal expression during disease progression.

## DISCUSSION

Plants respond to pathogen infection in a heterogeneous manner. Here, we revealed heterogeneity of plant responses at single-cell resolution using scRNA-seq coupled with confocal imaging of transcriptional reporter lines. Individual plant cells at immune, susceptible or transition states highlighted the gradient of responses within an infected leaf. Immune markers exhibit diverse spatial and temporal expression patterns while susceptible markers exhibit more expansive and sustained expression patterns in response to virulent *Pst* DC3000 (Figure 6). These data indicate that virulent bacteria are able to reprogram larger sections of the leaf towards susceptibility.

**Figure 6.**
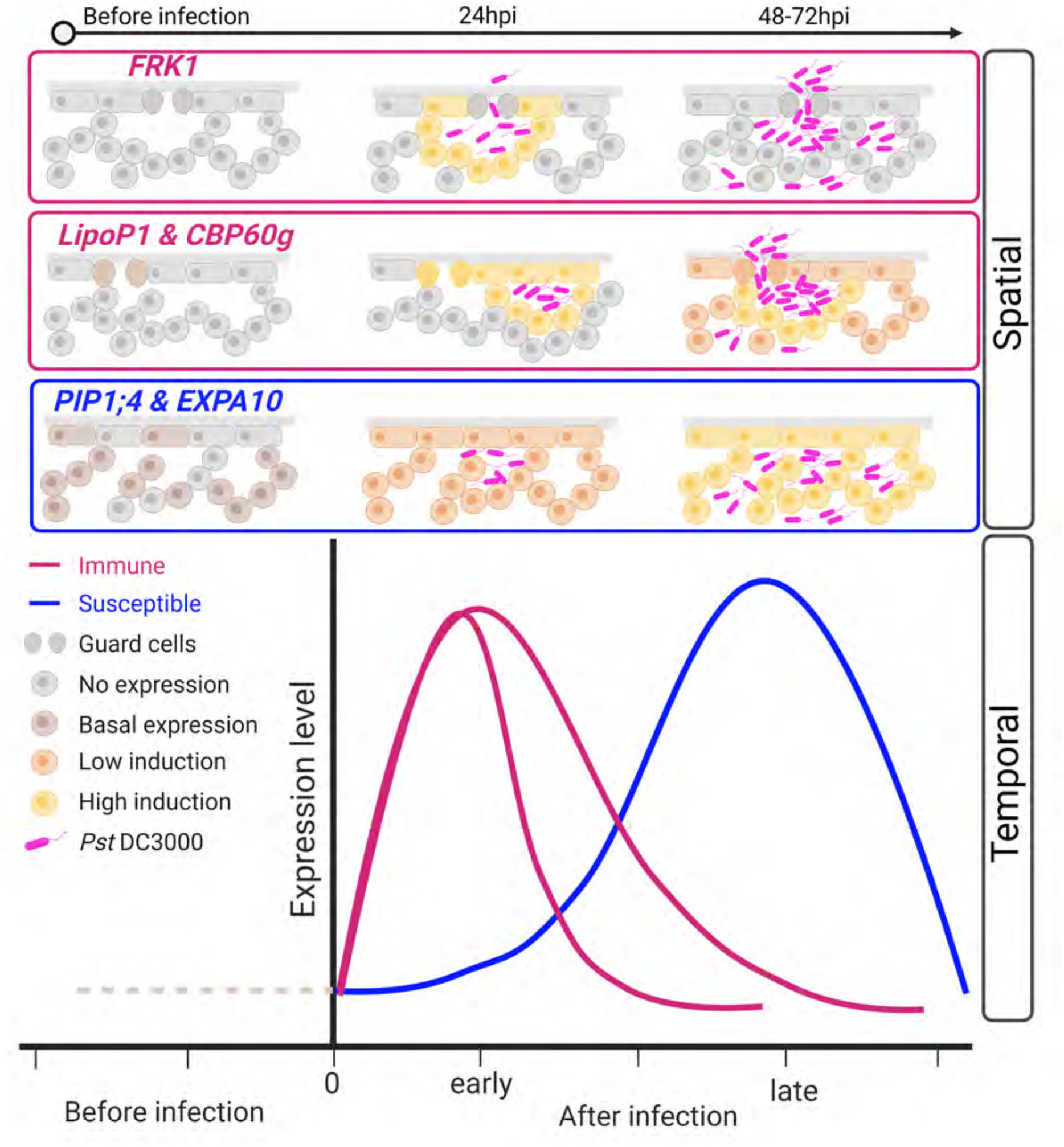
Spatial and temporal model of immune and susceptible marker gene expression. Immune and susceptible cell cluster markers exhibit diverse spatial and temporal expression patterns. Immune markers are highly induced at early infection stages, with in close proximity to bacterial colonies. At later time points (after 48h), the *FRK1* marker is no longer expressed, while other immune markers including *LipoP1* and *CBP60g* exhibit bundled and marginal expression patterns. Susceptible cell cluster markers exhibit basal expression that is broadly induced after infection throughout the leaf and peaks at late stages. Created with BioRender.com

Our understanding of host-pathogen interactions is largely influenced by assays investigating whole tissue samples. However, even after uniform inoculation, pathogens exhibit uneven penetration into the leaf interior and variable colonization within a tissue, which should result in variable host responses. For example, spores from the fungal pathogen *Zymoseptoria tritici* are able to continuously germinate on wheat leaves and their hyphae penetrate stomata for up to 10 days, resulting in multiple asynchronized infection stages at any given time (Fantozzi et al., 2021; Fones et al., 2017; Haueisen et al., 2019). *P. syringae* also exhibits uneven distribution on bean leaf surfaces, forming aggregates at leaf veins, crevices, trichomes and occasionally stomata (Björklöf et al., 2000). Similarly, we were able to visualize uneven colonization patterns of *Pst* DC3000 after syringe and surface inoculation on *Arabidopsis* (Figure 1A and 3A). Our sc-RNAseq analyses were able to simultaneously identify cell clusters exhibiting opposing biological processes (immunity and susceptibility) at 24h post-inoculation, indicating asynchronous infection stages even early during infection (Figure 2).

Although *Pst* DC3000 is virulent on *Arabidopsis*, the pathogen can still induce damage and carries MAMPs that can be perceived by plant pattern recognition receptors (flagellin, elongation factor Tu and 3’OH fatty acid epitopes), resulting in localized immune activated cell clusters. Previous research has found purified MAMP treatment can initiate transcriptional responses within 5 min and *Pst* DC3000 infiltration within 2h (Bjornson et al., 2021; Lewis et al., 2015). The pseudotime trajectory of our scRNA-seq data placed the immunity cell clusters at an early stage of disease progression (Figure 2D). We identified two immune cell clusters (Figure 2), possibly due to waves of PTI transcriptional responses (Bjornson et al., 2021). Consistent with these observations, our immune transcriptional reporter lines were also highly expressed at early infection time points (Figure 3). *FRK1* is a well-known early marker gene of PTI (He et al., 2006; Lewis et al., 2015; Zhou et al., 2020). We found *FRK1* expression was activated in cells surrounding substomatal cavities colonized by *Pst* DC3000 (Figure 3B and 5A-C). *Pst* DC3000 uses stomatal pores to enter leaves and substomatal cavities are an early site of pathogen colonization (Melotto et al., 2008). Similarly, the highly induced immune markers *LipoP1* and *CBP60g* exhibited proximal expression to bacterial colonies during infection (Figure 5D-H and Figure S7C-E). The clustering of immune markers in cells surrounding bacteria may indicate that these bacterial colonies carry or create sufficient MAMPs/DAMPs to induce defense. Compared to expression surrounding bacterial colonies, the immune marker *CBP60g* exhibited weaker, but general induction, in most cells at all infection stages, consistent with its role in inducing SA synthesis whose accumulation is required for defense within a leaf as well as systemic immune responses (Figure 5G) (Kim et al., 2022; Sun et al., 2015).

Plant pathogens deliver effectors into host cells to dampen immune responses and promote susceptibility (Garcia-Ruiz et al., 2021; Toruño et al., 2016). The timing and number of cells targeted for effector delivery also varies, but can occur within 75-90 minutes after bacterial infiltration on *Arabidopsis* (Grant et al., 2000; Henry et al., 2017). Transcriptional profiling of *Arabidopsis* infected by virulent *Pst* DC3000 detected effector-mediated suppression of PTI and upregulation of genes contributing to susceptibility by 6h (Lewis et al., 2015; Nobori et al., 2018). Our scRNA-seq and promoter reporter line investigations identified clusters of cells exhibiting patterns consistent with a compatible or susceptible interaction that peaked later during infection (Figure 2 and 4). For example, genes involved in water transport and abscisic acid (ABA)related processes were enriched in susceptible (M4-M5) clusters. Recently, it has been reported that the HopM1 and AvrE effector family are responsible for apoplastic water-soaking phenotypes as a result of their ability to manipulate ABA signaling to induce stomatal closure (Gentzel et al., 2022; Hu et al., 2022; Roussin-Léveillée et al., 2022). Unlike the localized expression of immune markers, susceptibility markers exhibited more general expression, indicating more global reprogramming of the leaf to a susceptible state over time. The susceptibility markers *EXPA10* and *PIP1;4* encode members of gene families that function in cell wall enlargement/expansion and water transport under normal conditions, respectively (Cosgrove, 2015; Maurel et al., 2015). During infection these processes can be manipulated by pathogens to create favorable environments for proliferation and disease development (Ding et al., 2008; Xin et al., 2016; Gentzel et al., 2022; Hu et al., 2022; Roussin-Léveillée et al., 2022).

Transcriptional profiling of entire tissues can mask cells at opposing response trajectories by averaging signals across thousands of cells. Comparing gene regulation in immune (M1-M2) and susceptible (M4-M5) clusters has resulted in the identification of novel candidates (Figure S4-S5 and Table S3), including specific members of large gene families, involved in foliar plant-pathogen interactions. Genes regulating plant immune perception and signaling have been well characterized over the past 30 years, but identification of susceptibility genes has lagged behind (van Schie and Takken, 2014). Susceptibility genes are attractive targets to modify for developing disease resistant crops due to advances in genome editing technologies and decreased regulatory oversight (Garcia-Ruiz et al., 2021; Zaidi et al., 2018). Future advancements in high-resolution spatial transcriptomics enabling profiling of both plant and pathogen tissues, will facilitate investigating gene expression in a positional context in complex tissues (Nobori et al., 2022; Saarenpää et al., 2022). Detailed characterization of cellular states throughout disease development will enable a comprehensive understanding of mechanisms regulating disease progression.

## ACKNOWLEDGMENTS

We thank Dr. Niko Geldner (University of Lausanne, Switzerland) for providing *FRK1* transcriptional reporter seeds, Dr. Lei Lei (Lanzhou University, China) for bulk RNA extraction from leaves and Dr. Pamela Ronald for use of their confocal microscope. We also would like to thank all the members in Coaker Lab for helpful discussions and input on the project. This work was supported by the grants to G. C. from the NIH R35GM136402. S.L. was supported by the Independent research fund Denmark (grant number 7026-00053B). B.K. was supported by a USDA NIFA Award 2017-67030-25920. The sequencing was carried out at the UC Davis Genome Center DNA Technologies and Expression Analysis Core, supported by a NIH Shared Instrumentation Grant 1S10OD010786-01.

## AUTHOR CONTRIBUTIONS

J.Z., S.L., B.C. and G.C designed the study. S.L. prepared samples for RNA-seq, B.C. analyzed sequencing data, J.Z. performed all microscopy and promoter-reporter analyses, A.T. and B.G. assisted with developing promoter-reporter lines. B.K. developed tagged *Pst* DC300. J.Z., S.L., B.K., B.C., and G.C. participated in the discussion and interpretation of results. J.Z, B.C., and G.C. wrote the manuscript. All authors reviewed and approved the manuscript.

## DECLARATION OF INTERESTS

The authors declare no competing interests

## STAR METHODS

## RESOURCE AVAILABILITY

### Lead contact

Further information and requests for resources and reagents should be directed to and will be fulfilled by the lead contact, Gitta Coaker

### Materials availability

Seeds of transgenic plants generated in this study are deposited in *Arabidopsis* Biological Research Center (ABRC, stock number: see key resource table). Plasmids used to generate transgenic plants are deposited in Addgene (192521-192525).

### Data and code availability

scRNA-seq and bulk RNA-seq datasets generated in this study have been deposited at Gene Expression Omnibus (GSE213625) and are publicly available as of the date of publication. Code describing data analysis performed in this study are available as a github respository (https://www.github.com/b-coli/AtPst_SingleCell). Original microscopy images have been deposited in Zenodo (https://doi.org/10.5281/zenodo.7686553) and are publicly available as of the date of publication.

## EXPERIMENTAL MODEL AND SUBJECT DETAILS

### Plant material and growth conditions

*Arabidopsis thaliana* ecotype Columbia Col-0 was used in single-cell RNA sequencing, bacterial growth curves and plant transformation. The transcriptional reporter line *pFRK1::NLS-3xmVENUS* in the Col-0 background was obtained from Professor Niko Geldner’s lab (Zhou et al., 2020).

*Arabidopsis thaliana* seeds (Col-0 or transgenic lines) were stratified for 2 days in the dark at 4°C before sowing onto soil or half-strength (1/2) Murashige and Skoog (MS) medium. Seeds were also surface-sterilized with disinfection solution (50% Bleach, 0.1% Tween-20) for 8 min and 75% ethanol for 1 min, washed thoroughly in sterile water for 4 times before sowing onto 1/2 MS medium. Four-week-old *A. thaliana* Col-0 used for scRNA-seq and plant transformation were grown in a controlled environment chamber at 22°C and 70% relative humidity with 10-h light/ 14-h dark photoperiod (100 μM m^−2^ s^−1^). Ten to 14-day-old seedlings grown on 1/2 MS were incubated at 22°C under long-day conditions with 16-h light/ 8-h dark cycles. Seedlings were used for microscopy analyses.

### Bacterial strains and growth conditions

*Pseudomonas syringae pv. tomato* DC3000 *ΔhopQ1* and the wilt-type *Pst* DC3000 were labeled with 3xmCherry (*att*Tn*7*-3xmCherry, *Pst* DC3000-mCherry) using the same approach described previously (Baltrus et al., 2022). Bacteria were cultured overnight at 28°C on nutrient yeast glycerol agar (NYGA) medium containing 100 µg/mL of rifampicin and 50 µg/mL of spectinomycin prior to inoculation.

## METHOD DETAILS

### Bacterial inoculation and quantification

Cells from an overnight culture of *Pst* DC3000-3xmCherry or *Pst* DC3000 *ΔhopQ1*-3xmCherry were collected and resuspended in 10mM MgCl_2_. For scRNA-seq, bulk protoplast RNA-seq, leaf RNA-seq samples, and bacterial growth curves, leaves of four-week-old *A. thaliana* were syringe infiltrated with a bacterial suspension of OD_600_ = 0.0001. Inoculated plants were kept under ambient humidity for 1 h to allow evaporation of excess water on the leaf surface. Then plants were covered with a transparent dome to maintain high humidity and incubated in growth chamber for 24h. For seedling flood inoculation, two-week-old plants on ½ MS medium were flood inoculated using 40 mL of the bacterial suspension of OD_600_ = 0.01 with 0.02% Silwet L-77 per 100 mm x 100 mm square petri dish (Fisherbrand). The bacterial suspension was removed after 20-30s incubation at room temperature. Inoculated plants were sealed with 3 M Micropore tape (3 M, St. Paul, MN, U.S.A.) and incubated in a growth chamber.

Bacterial titers after flood inoculation were determined as colony-forming units (CFU) per milligram. In brief, three plants were cut roots away as one biological repeat, and 5-7 repeats were taken for each time. After measuring the weight of the aerial parts of each repeat, plants were ground and diluted in 5mM MgCl_2_. The bacterial suspensions were then dilution plated on (NYGA) medium containing 100 µg/mL of rifampicin. Colonies were counted at each time point after incubation at 28°C. Bacterial titers after syringe infiltration were determined as described previously (Liu et al., 2009)

### Protoplast isolation

Protoplasts were isolated from *Arabidopsis* leaves infiltrated by bacteria and 10 mM MgCl_2_ (Mock) using Tape-*Arabidopsis* Sandwich method as described previously (Wu et al., 2009). The adaxial side of 10-20 infiltrated leaves for each treatment was stabilized on the time tape and the abaxial side was adhered to the Magic tape (3M). The abaxial side was removed by carefully pulling off the Magic tape. Peeled leaves were immediately immersed in a petri dish containing 10 mL of enzyme solution (1.5% Cellulase Onuzuka R-10 (Yakult, Japan), 0.3% Macerozyme R-10, (Yakult, Japan), 0.4 M Mannitol, 20 mM KCl, 20 mM MES (2-(N-Morpholino)ethanesulfonic acid hydrate) pH 5.7, 10 mM CaCl2, and 0.1%, BSA). After digesting for 100 min with gentle shaking, the protoplast suspension was filtered through 40 μm cell strainer (BD Falcon 352340) into a round-bottomed 50 mL tube and centrifuged at 100 x g for 1 min at 22°C using a swinging rotor. Protoplast pellets were gently resuspended in 10 mL of CS-sucrose buffer (0.4 M sucrose, 20 mM MES pH 5.7, 20 mM KCl) and centrifuged at 100 x g for 2 min at 22°C. Intact and healthy protoplasts remained suspended in the upper layer. The upper layer suspension was then transferred into a clean round-bottomed tube and gently mixed with 10 mL of protoplast buffer (0.4 M Mannitol, 20 mM KCl, 20 mM MES pH 5.7, and 0.1%, BSA). After centrifuging at 100x g for 2 min at 4°C using a swinging rotor, the supernatant was removed without disturbing the loosely packed protoplast pellets. The protoplast concentration was determined using a hemocytometer and the viability was checked using trypan blue solution. The protoplast sample from each treatment (DC3000 and mock) was divided for scRNA-seq and bulk RNA-seq.

### scRNA-seq library preparation and sequencing

The protoplast suspension was diluted to a final concentration of 1000 cells/µL. A total of 40, 000 cells were loaded into a microfluidic chip (10X Genomics) with v3 chemistry to capture ∼10,000 cells per sample. Protoplasts were barcoded with a Chromium Controller (10X Genomics). mRNA was reverse transcribed and cDNA libraries were constructed with a Chromium Single Cell 3’ reagent kit V3 (10X Genomics) according to the manufacturer’s instructions. Eleven cycles were used for cDNA amplification and 10 cycles were used for final library amplification. cDNA and final library quality was assessed using a Bioanalyzer 2100 High Sensitivity DNA Chip (Agilent). Sequencing of paired-end 150 bp reads was performed with a NovaSeq 6000 instrument (Illumina) at the University of California Davis Genome Center. Protoplasts from DC3000 treatment and mock inoculation were each barcoded on a single Chromium Controller.

### RNA extraction, bulk RNA-seq library preparation and sequencing

Total RNA was extracted with TRIzol (Fisher #15596018), following the manufacturer’s instructions, for infiltrated leaves and leaf protoplasts isolated using the above mentioned method for scRNA-seq. DNase treatments were performed with RQ1 RNase-Free DNase (Promega #PR-M6101). Three biological replicates were performed for samples of *Pst* DC3000-or mock-infiltrated leaves, and one repeat was made for leaf protoplasts of each sample. cDNA libraries were prepared with QuantSeq FWD kit (Lexogen), according to the manufacturer’s protocol. The fragment size distribution was evaluated by a Bioanalyzer 2100 (Agilent). The library pool was treated using Exonuclease VII (NEB), SPRI-bead purified with KapaPure beads (Kapa Biosystems/Roche), quantified via qPCR with a Kapa Library Quant kit (Kapa Biosystems) on a QuantStudio 5 RT-PCR system (Applied Biosystems). Sequencing was performed at the University of California Davis Genome Center using a HiSeq 4000 (Illumina) platform with single-end 100-bp reads.

### Bulk RNA-seq data analysis

Raw fastq files for three bulk RNA-seq replicates each for *Pst* DC3000- and mock-inoculated leaves, as well as one bulk RNA-seq replicate from protoplasts isolated from *Pst* DC3000- and mock-inoculated leaves were trimmed using TrimGalore (stringency = 4, default parameters otherwise) and aligned to the *Arabidopsis thaliana* reference genome (Araport11) and quantified using STAR. Differential expression metrics for protoplasting and bacterial-induced changes were evaluated using the glmQLFit method from the edgeR package (Bioconductor v3.12), using a design matrix that takes into consideration the interaction between these two variables. Genes determined to be significantly altered by protoplasting were identified using a relatively liberal set of criteria, i.e. having log-fold change values > 0.5 and adjusted (BH) p-values less than 0.05, and were removed from dimension reduction and integration analyses. Genes determined to be significantly affected by *Pst* DC3000 were those having log-fold change values >2 and adjusted (BH) p-values less than 0.01, unless otherwise specified.

### scRNA-seq data initial processing and integration

Raw fastq files for the two samples generated in this study (Mock, *Pst* DC3000) were processed using Cellranger (v6.0.1; 10x Genomics, Pleasanton, CA) using default parameters (and an expected cell # equal to 10,000), mapping to the Araport11 *Arabidopsis* reference genome. The output from Cellranger was further processed using the Velocyto (v0.17.15) algorithm (La Manno et al., 2018), using default parameters, to generate spliced and unspliced counts matrices. For each cell, the percentage of reads mapping to mitochondrial, and chloroplast genes was computed. Cells were then filtered for those having a spliced mitochondrial read percentage of less than 1%, as well as a total spliced Unique Molecular Identifier (UMI) count within a dataset dependent threshold, bounded at the high end by 50,000 counts, and at the low end by 10% of the UMI count of the 100^th^ most spliced transcript-rich cell for that dataset (Zheng et al., 2017).

Cells were normalized using the SCTransform method (Seurat, v3.9.9005). A recent *Arabidopsis* leaf single-cell RNA-seq dataset (Kim et al., 2021) was used to annotate cell types for all cells in this dataset using the label transfer pipeline (Seurat). Genes that were identified as being significantly influenced by protoplasting (see Bulk RNA-seq data analysis) were excluded from further analysis.

The *Pst* DC3000 and mock-inoculated datasets were then integrated using the anchor method (Seurat). Fifty principal components were calculated for the integrated dataset, used to cluster the cells (Louvain method, resolution 0.8) and further dimensionally reduce the gene expression space using Uniform Manifold Approximation and Projection (Seurat), using 50 Principal Components and default parameters (Table S6).

### Pseudobulk analysis

A pseudobulk value for each gene was calculated as the sum of all counts from all cells for that gene within either the Mock- or *Pst* 3000-treated single-cell datasets. These values were then used to compare against whole-tissue or pooled-protoplast bulk RNA-seq data to verify that the single-cell datasets coarsely resemble bulk RNA-seq datasets.

### Signature score computation

A *Pst* DC3000 signature score was computed as a composite metric quantifying the overall impact that *P. syringae* has on each cell. Here, the Seurat AddModuleScore function was used to define a pair of module score for genes up- or down-regulated by *Pst* DC3000 (from bulk RNA-seq data), with the signature score defined as the *Pst* DC3000-up module score subtracted from the *Pst* DC3000-down module score. Similarly, an Immunity Response Score was defined as a composite signature score quantifying the general state that each cell was in with respect to disease progression based on sets of genes known to be induced/involved in immune response (Immunity) and in advanced disease (Susceptibility),. These genes were filtered for those that were found to be differentially expressed from our bulk RNA-seq analysis (see above). Immune and Susceptibility module scores were then computed using the AddModuleScore function (Seurat), and the Response Score was defined as the Susceptibility score subtracted from the Immunity score. Similarly, we also generated a protoplast signature score for those genes induced or repressed by protoplasting, generating another compound signature score for the overall effect of protoplasting (Figure S2)

### Cluster-specific marker loci

Marker genes specifically expressed in each cell cluster were determined using the FindAllMarkers function (Seurat) (Table S4). Significant markers were defined as those having a log-fold change (compared to all other clusters) greater than 0.25, and an adjusted (BH) p-value (wilcoxon rank sum test) less than 0.01. Log-fold change values between Mock- and *Pst* 3000-treated single-cell transcriptomes were computed for all superclusters (Immunity, Susceptibility, Transition, etc.) using the FindMarkers function (Seurat), with significance calculated using the DESeq2 (v1.30.1) method on the unnormalized counts.

### GO Term Enrichment

For each cluster, a stringent set of marker loci was computed using the FindAllMarkers function in Seurat. GO term enrichment analysis was then performed using the topGO R package (Bioconductor version 3.12) for these marker genes, using all expressed genes (excluding those induced by protoplasting) as background. GO enrichment was calculated as the number of significant genes divided by the number of expected genes for each GO term.

### Pseudotime inference

SCTransform-normalized expression values for spliced transcripts in mesophyll cells (excluding Seurat cluster 16, which seemed distinct from other mesophyll cells) were filtered from the *Pst* DC3000 dataset and re-embedded in a low-dimensional UMAP space using the Monocle3 (v1.0.0) pipeline (using 5 principal components, the correlation distance metric, and a minimum distance of 0.01). A cell trajectory was then imputed using Monocle3, defining the starting cell as that with the lowest *Pst* DC3000-expression score (a measure of how influenced the cell is by pathogen expression, empirically determined using bulk RNA-seq expression data) within the cluster with the lowest mean *Pst* DC3000 signature score, and a minimum branch length of 15. Pseudotime was projected onto this cell trajectory for *Pst* DC3000 mesophyll cells. Genes that vary significantly with pseudotime were computed with the graph_test function (Monocle3), using “principal graph” as the neighbor_graph parameter. Genes were selected as significant as those having a Morans I value greater than 0.2 (Table S3).

### Generation of transgenic lines

Genes for the generation of reporter lines were selected based on their enrichment in clusters M1-M5 (adjusted p-value from the FindAllMarkers Seurat function less than 0.01, and a log-fold change greater than 1), relatively specific expression from scRNA-seq analyses in either immune or susceptible clusters and potentially interesting functions from the literature. Promoters (∼ 2 kb upstream of the start codon) of *LipoP1* (AT3G18250, 2210 bp), *CBP60g* (AT5G26920, 2183 bp), *EXPA10* (AT1G26770, 2025 bp), *PIP1;4* (AT4G00430, 2151 bp) and *ILL5* (AT1G51780, 2089 bp) were PCR-amplified and fused to a nuclear localization signal (NLS) in pENTR vectors using In-Fusion HD Cloning Plus (Clontech). See Table S5 for primer details. The resulting constructs were recombined with binary destination vector pMpGWB123 (Ishizaki et al., 2015) bearing 3xmCitrine using Gateway Cloning Technology (Invitrogen). All plasmids were transformed into *Agrobacterium tumefaciens* GV3101 strain and then transformed into *A. thaliana* Col-0 by floral dipping method (Zhang et al., 2006). Seedlings of transgenic plants were screened on ½ MS plates supplemented with 25 µg/mL of hygromycin and 100 µg/mL of carbenicillin. Ten to fifteen independent T1 lines were analyzed for mCitrine fluorescence, and 3-5 T2 lines expressing mCitrine were selected for bacterial infection. Transgenic plants T2 or T3 (1-2 independent lines) with similar induction patterns after bacterial infection were selected for further experiments.

### Confocal settings and image processing

Confocal imaging was performed on either a Leica TCS SP8 or Zeiss LSM 980 with Airyscan 2 laser scanning microscope. Pictures were taken with a 20x (Leica TCS SP8 or Zeiss LSM 980), 10x dry immersion objectives (Leica TCS SP8), as well as 5x immersion objective for tile-scan with 10% overlap (Leica TCS SP8). The following excitation and emission parameters were used for different fluorophores: mVENUS/mCitrine 488 nm, 493 – 540 nm; mCherry 552 nm, 586 – 635 nm; chlorophyll 638 nm, 650 -720 nm on Leica TCS SP8. mCitrine 488 nm, 490 – 543 nm; mCherry 561nm, 570 – 640 nm on Zeiss LSM 980. Sequential scanning was applied to avoid fluorescence interference between channels. Time-course confocal images of each transcriptional reporter line were taken under identical settings (lens, laser power, pinhole size, detector gain and interval of Z stack) for comparison of fluorescence intensity over time. Different microscope settings were applied for different transcriptional reporter lines according to the expression level of transgenes in plants.

Quantification of bacterial colony number, area, and fluorescence intensity was performed with the Imaris software (https://imaris.oxinst.com/). In brief, the Surface Model tool was used to quantify colony number and area of an entire image.

Background subtraction was used to manually adjust threshold until all visible fluorescently-labeled bacteria colonies were detected. Then the detected colonies were counted and surface area measured. The Spot tool was used to build spots for each fluorescence domain to quantify fluorescence intensity. Quantification was determined by measurement of mean fluorescence intensity per spot (fluorescence intensity/nucleus). Algorithm settings of Different Spot Sizes were used due to variable nuclei size in different cell types of plant leaves. Background subtraction function was used for spot detection. To classify spots, “Quality (pixel intensity of a spot center)” filter was used to manually adjust threshold until all visible fluorescently-labeled nuclei were detected. Spot regions were determined by manually adjusting threshold of absolute intensity detection to ensure complete coverage of fluorescently-labeled nuclei.

## STATISTICAL ANALYSIS

Statistical analyses were performed with Graphpad Prism 9.0 software (https://www.graphpad.com/) or in R. The data are presented as mean ± SD, and “n” represents number of analyzed images from at least 3 plants. One-way ANOVA with Tukey’s test was used for multiple comparisons. Two tailed Student’s t-test was used to compare means for two groups. Details about the statistical analyses are described in the figure legends.

## SUPPLEMENTAL FIGURE TITLES AND LEGENDS

**Figure S1.**
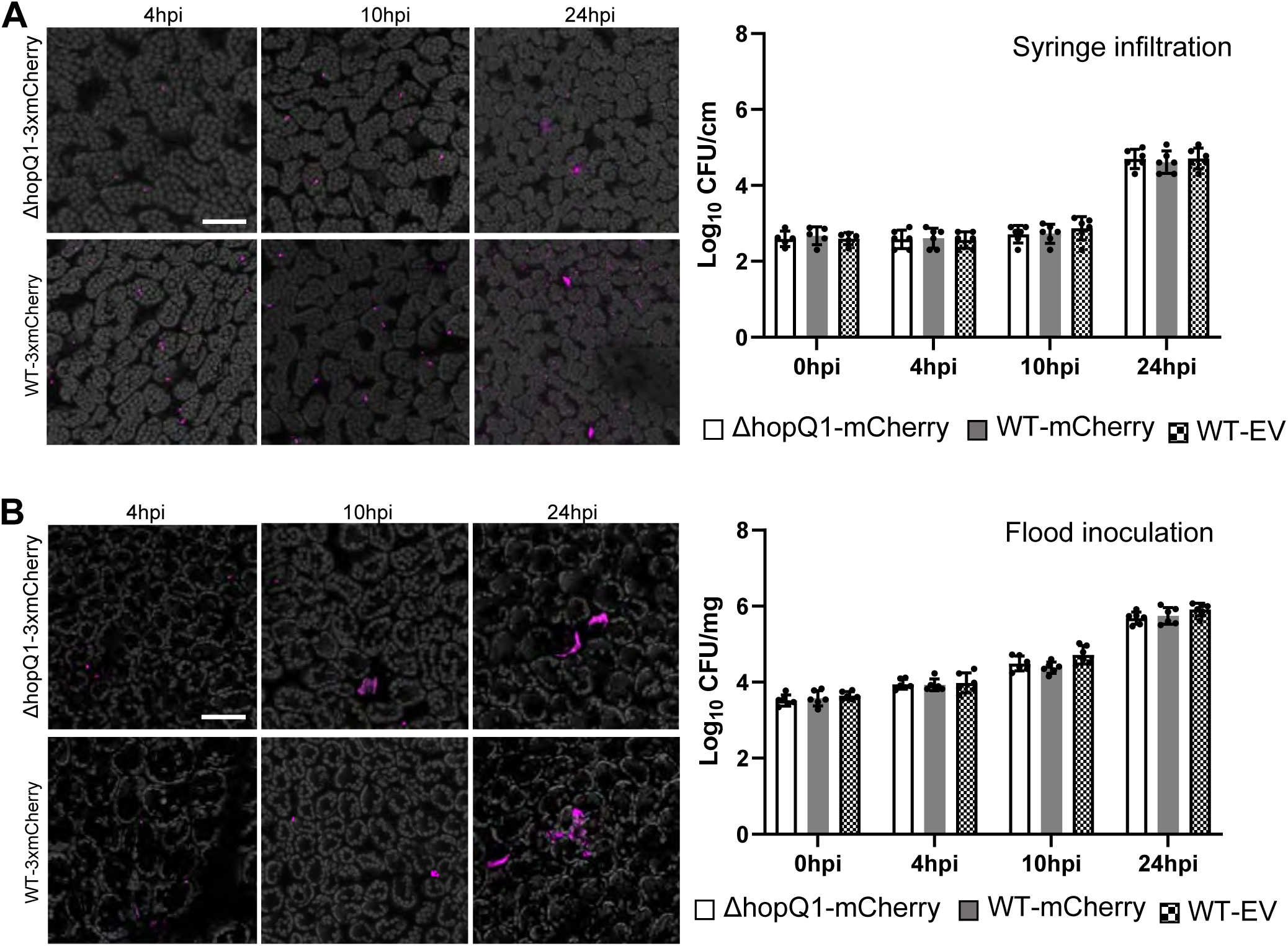
Comparison of *Pst DC3000 WT-3xmCherry* and *Pst DC3000 ΔhopQ1-3xmCherry* during infection in *Arabidopsis*. Related to Figure 1. (A) Bacterial populations in wild type *Arabidopsis* Col-0 leaves using syringe infiltration. Four-week-old *A. thaliana* leaves were syringe infiltrated with a bacterial suspension of OD_600_ = 0.0002. Left: Representative images of bacterial colonization at different time points. Pictures are maximum projections from confocal Z stacks. Right: Bacterial growth curve over time. Log_10_ CFU/cm^2^, log_10_ colony forming units per cm^2^ of leaf tissue. Data are means ± SD (at least 5 plants used for each strain at each time point). (B) Bacterial populations in wild type *Arabidopsis* Col-0 leaves using flood inoculation. Two-week-old *Arabidopsis* seedlings grown on Murashige-Skoog plates were flood-inoculated with mCherry-tagged *Pst* DC3000 at concentration of 1 x 10^7^ colony forming units/ml (CFU/ml). Left: Representative images of bacterial colonization at different time points. Right: Bacterial growth curve over time. Log_10_ CFU/mg, log_10_ colony-forming units per milligram leaf tissue. Data are means ± SD (at least 6 plants used for each strain at each time point). Pictures are maximum projections from confocal Z stacks. Chlorophyll autofluorescence is shown in gray. Scale bars: 50 µm.

**Figure S2.**
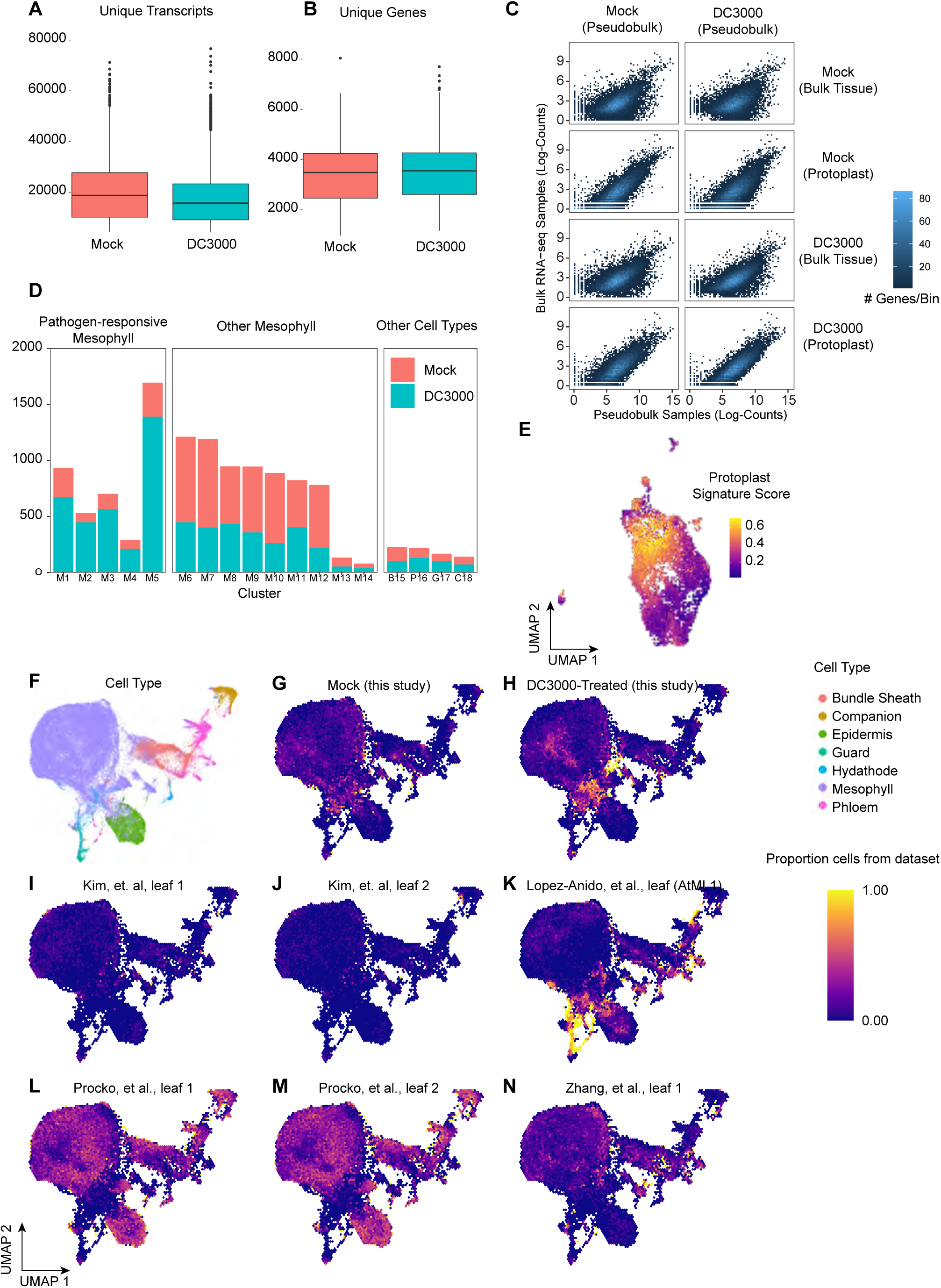
Data quality of single-cell datasets. Related to Figure 1-2. (A-B) Summary statistics of single-cell datasets. Transcript (A) and Gene (B) counts were summarized across all 11,895 cells in the integrated single-cell dataset. Median transcript counts were ∼ 17,000 unique molecular identifier (UMI) per cell for both datasets, while median gene counts were ∼ 3,500. (C) Comparison with bulk RNA-seq. For Mock- and DC3000-treated single-cell datasets, a “pseudobulk” profile was calculated as the sum of counts per gene for all cells within that dataset. These profiles were compared to mRNA-seq data derived from whole leaves (Bulk Tissue) and pooled protoplasts (Protoplast) isolated in the same manner as those used for single-cell profiling). (D) Cell numbers from *Arabidopsis* leaves infiltrated with *Pst* DC3000 or 10mM MgCl_2_ (Mock) in each cluster. (E) The protoplast signature score is depicted in the UMAP of the combined scRNA-seq datasets. A module score (Protoplast Signature Score) was computed based on genes up- and down-regulated by protoplasting (identified from bulk/protoplast RNA-seq). While heterogeneous, protoplasting does not appear to strongly impact cells within immune/transition/susceptibility clusters. (F-N) Integration of five leaf single-cell datasets from *Arabidopsis* preserves the unique identity of pathogen-treated cells. We integrated *Arabidopsis* leaf single-cell datasets (eight total, over five distinct studies: Kim et al., 2021; Liu et al., 2020; Lopez-Anido et al., 2021; Procko et al., 2022; Zhang et al., 2021) together to determine whether pathogen-treated cells remain distinct relative to other sources of cell-cell variation. (F) Cell type distribution in UMAP space for the integrated dataset. (G-N) Proportion of cells in UMAP space belonging to each individual dataset. While the combination of all datasets largely preserved clustering with respect to cell type (F), a region in UMAP space still appeared to be distinct for having *Pst* DC3000-treated identity (H). Heatmap represents proportion of cells for each dataset overlaid onto the integrated UMAP.

**Figure S3.**
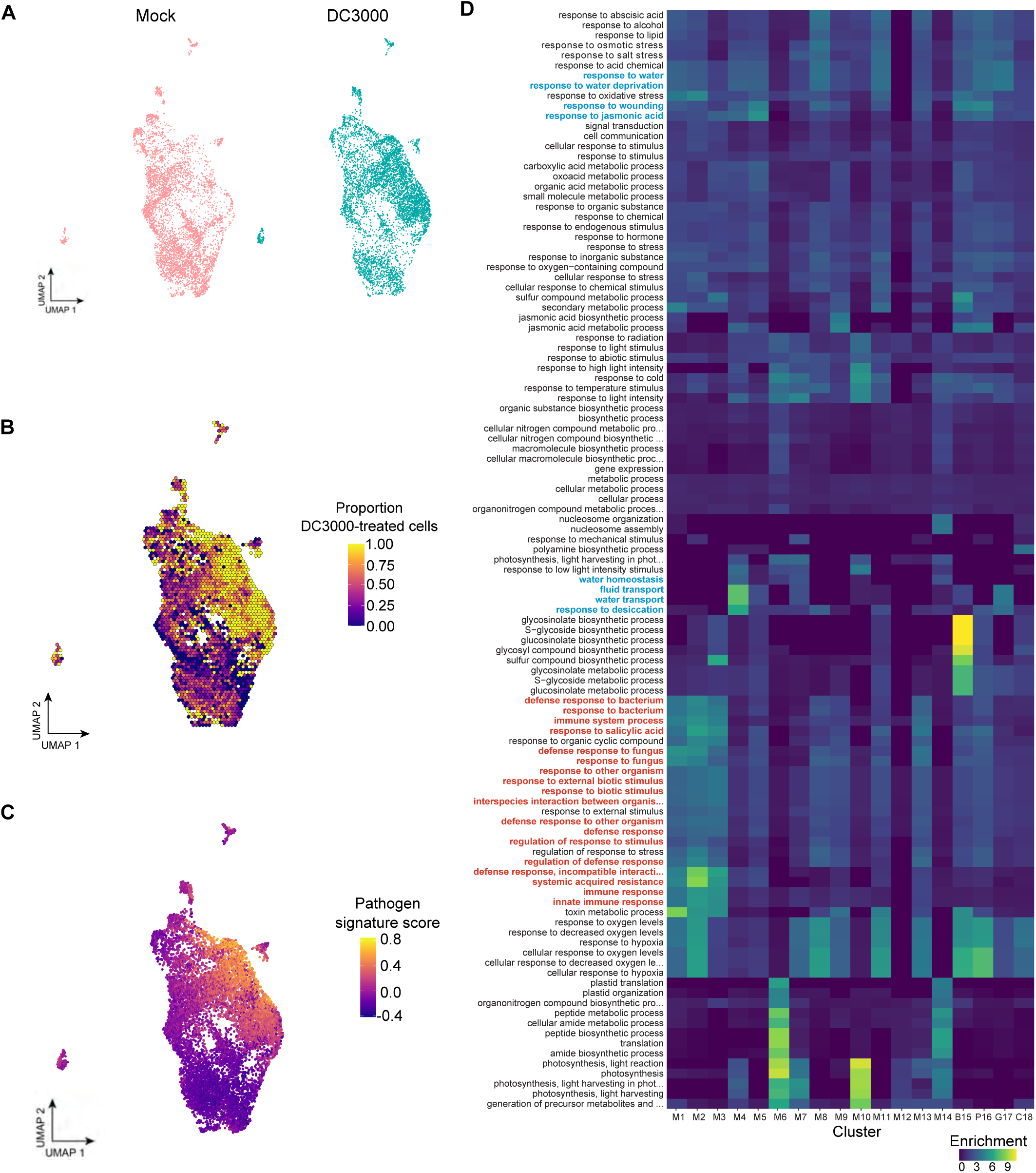
Clusters M1 – M5 are pathogen-responsive clusters. Related to Figure 2. (A) Single Cell Uniform Manifold Approximation and Projection (UMAP) plots from mock-treated samples (Mock) and *Pst* DC30000-treated samples, colored according to treatment. (B) Proportion of DC3000-treated cells is depicted in the UMAP of the combined single-cell RNA-seq datasets. These cells likely represent those that are responsive to pathogen infection. (C) A *Pst* DC3000 signature score was computed as a composite metric quantifying the overall impact that *P. syringae* has on each cell (Methods). The pathogen signature score is depicted in the UMAP of the combined scRNA-seq datasets. (D) GO-term enrichment (number of significant genes divided by the number of expected genes for each term) shows many terms specific to immunity and defense (bold, red) are associated with clusters M1-3, while terms specific to jasmonic acid, wounding and water stress (blue) are associated with clusters M4-5.

**Figure S4.**
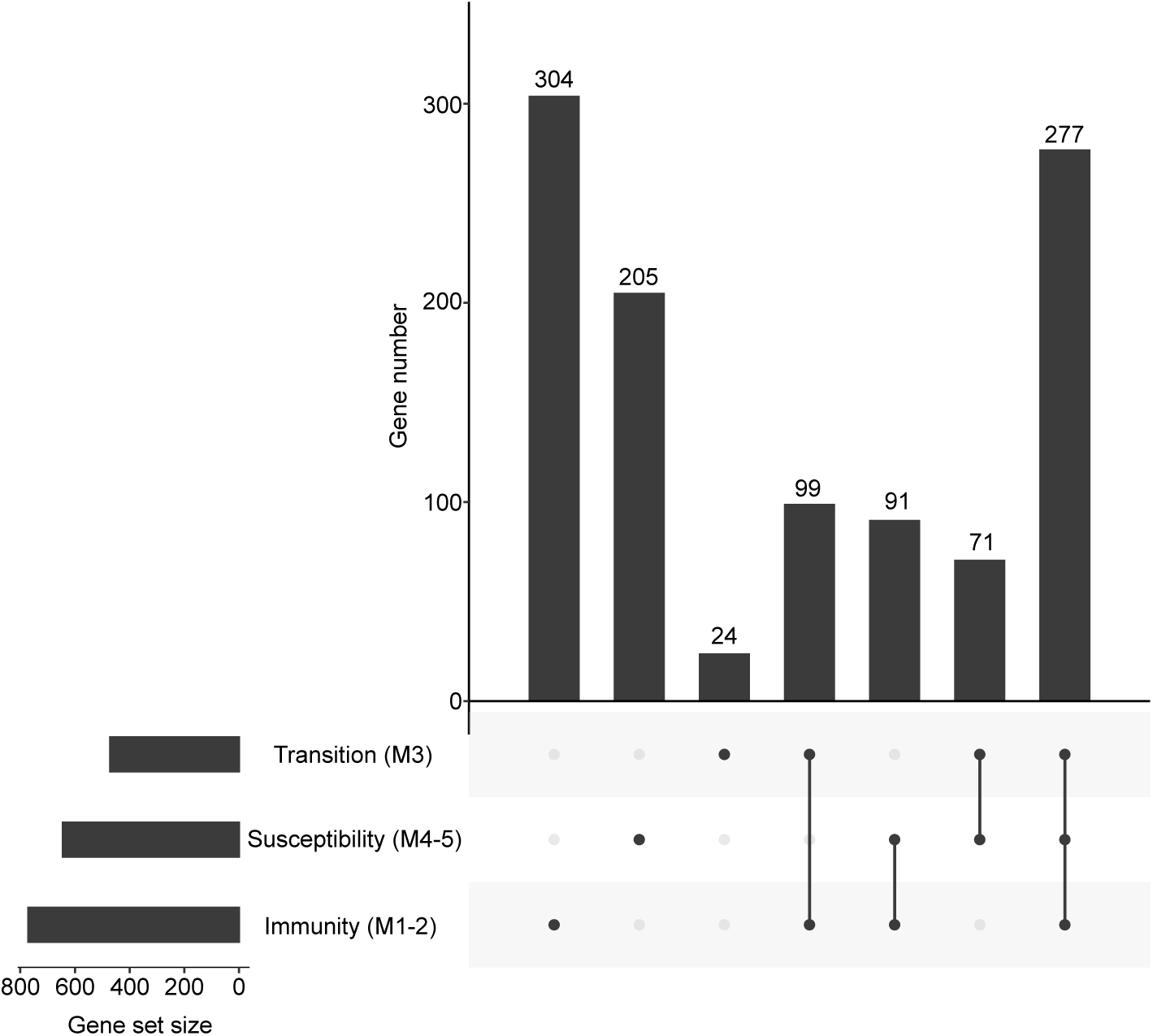
Overlap and specificity of differentially expressed genes in pathogen responsive clusters. Related to Figure 2. Overlap among genes identified as differentially expressed among different pathogen-responsive mesophyll cell types in single-cell data (log2-fold change > 0.5, adjusted p-value < 0.01, and negative binomial test using DESeq2).

**Figure S5.**
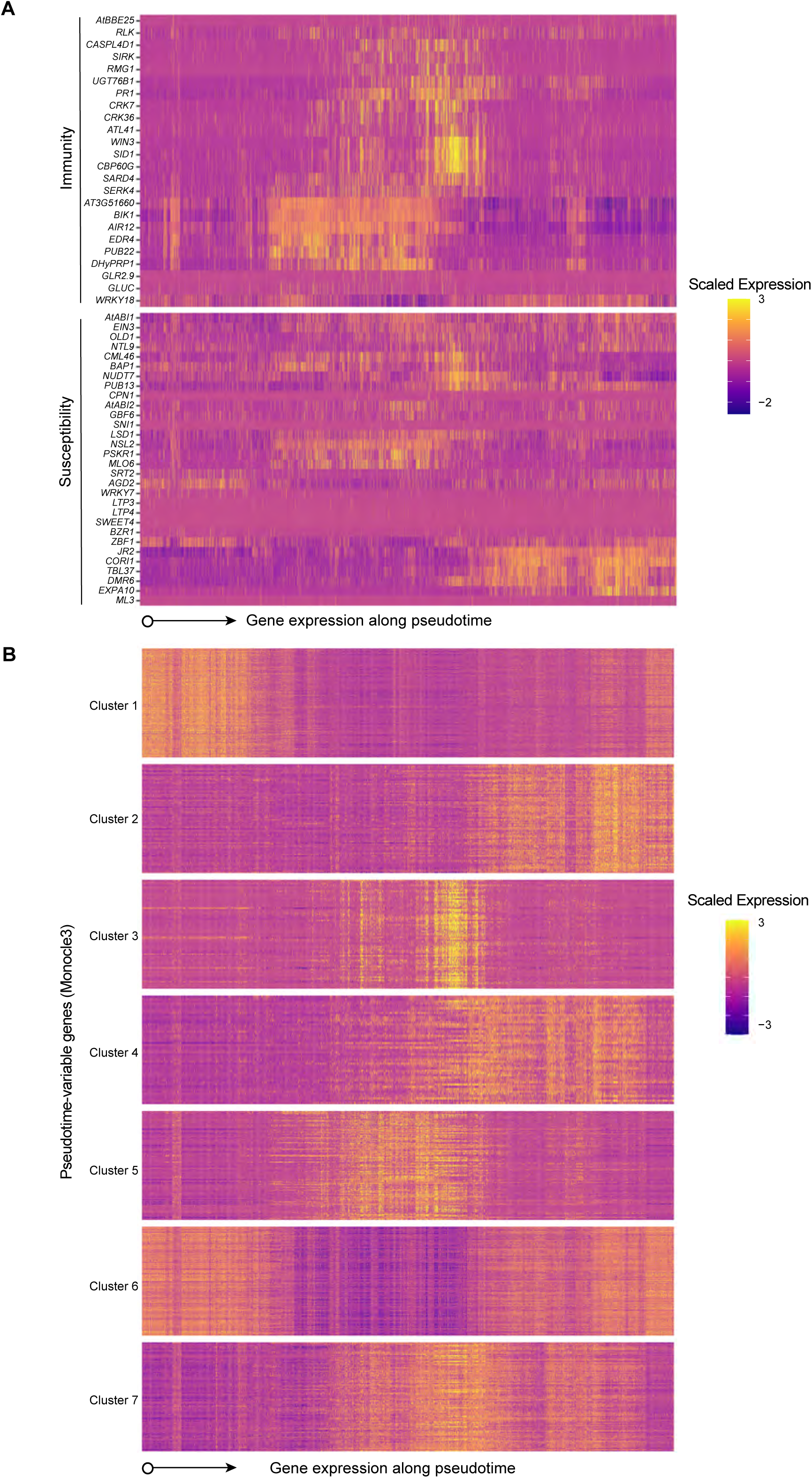
Variation of known and novel pathogen-responsive genes along pseudotime. Related to Figure 2 and Table S3. (A) Expression is shown for genes known to be involved in pathogen responsiveness. Known genes are organized based on their suspected involvement in immunity (top) or in disease susceptibility (bottom) and expression in individual cells are depicted varying along pseudotime on the x-axis. (B) Novel clusters of genes exhibiting similar expression patterns that vary with pseudotime were identified using monocle3. Novel genes are organized based on a hierarchical clustering of scaled expression profiles over pseudotime. Y-axis represents individual genes within a cluster, and the X-axis represents expression in individual cells organized along pseudotime.

**Figure S6.**
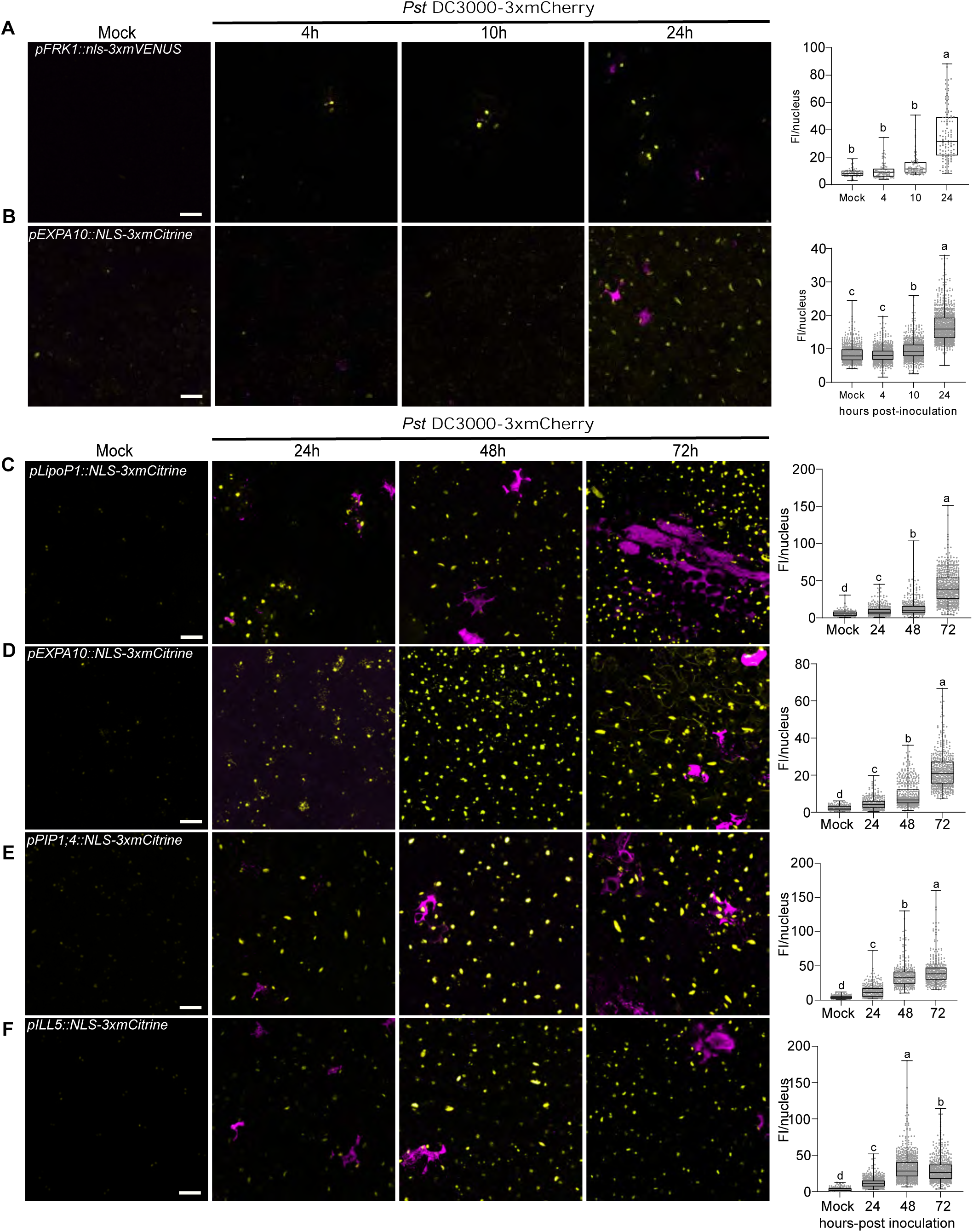
Visualization of immune and susceptible cellular markers during disease progression. Related to Figure 3-5. (A-B) Expression of immune marker *FRK1* and susceptible marker *EXPA10* at early infection stages. Two-week-old transgenic *Arabidopsis* seedlings grown on Murashige-Skoog plates were surface-inoculated with mCherry-tagged *Pst* DC3000 at concentration of 1 x 10^7^ colony forming units/ml (CFU/ml). Left: Representative images of marker gene expression at different infection stages. Mock images are taken at 24h. Pictures are maximum projections from confocal Z stacks. Right: Mean florescence intensity (FI, mean gray values) per nucleus was calculated and boxplot shows median with minimum and maximum values indicated (n = 6 images from 3 plants). Different letters indicate statistically significant differences (p < 0.0001, ANOVA with Tukey test). Scale bars: 50 µm. Experiments were repeated two times with similar results. (C-F) Data represent experiments with a second, independent transgenic line for promoter-reporter constructs (line 2). (C) The immune marker *LipoP1* is highly expressed during infection. The promoter-reporter line for this immune marker was generated with fusion to 3xfluorophore possessing a nuclear localization signal (NLS). Plants were inoculated as described in (A). (D, E and F) The susceptible markers *EXPA10* (D), *PIP1;4* (E) and *ILL5* (F) are highly induced at late infection stages. Promoter-reporter lines for each susceptible marker were generated as mentioned in (C).

**Figure S7.**
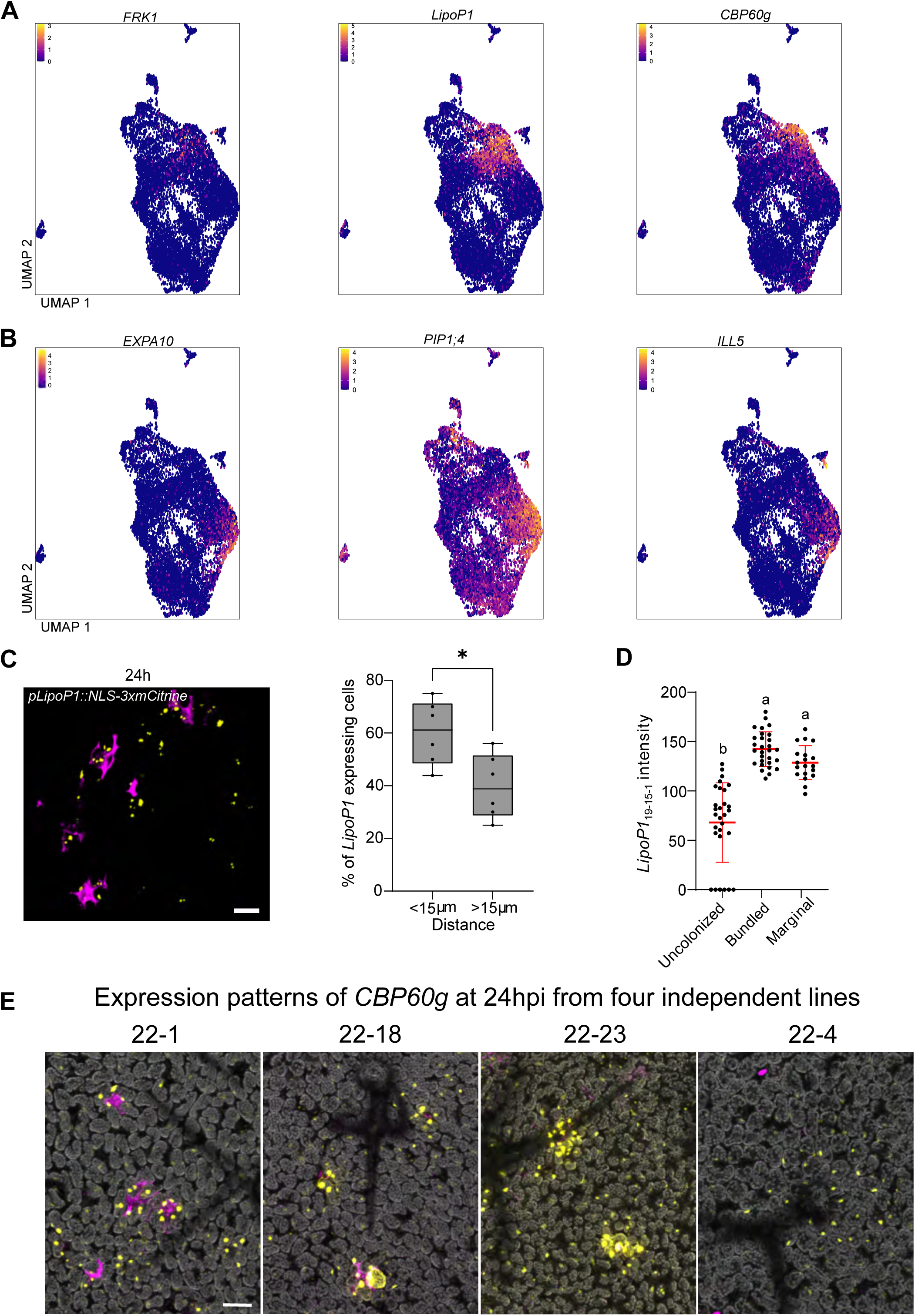
Feature plots of marker genes and expression pattern of immune markers. Related to Figure 3-5. (A-B) Feature plots of immune (A) and susceptible (B) markers in all cell clusters. (C) Expression of the immune marker *LipoP1* (line 19-15-1) is proximal to bacterial colonies. Two-week old *Arabidopsis pLipoP1::NLS-3xmCitrine* seedlings were flood-inoculated with mCherry-tagged *Pst* DC3000. Left: Maximum projections of Z stack of mVENUS and mCherry signals. Each yellow dot indicates a single nucleus. Scale bar: 20 µm. Right: Percentage of *LipoP1* expressing cells that are proximal (<15 µm) or distal (>15 µm) to a bacterial colony 24hpi. Boxplot shows median with minimum and maximum values indicated (n = 6 images from 3 plants). *p < 0.05 analyzed by two-tailed, unpaired Student’s t-test. (D) Expression of *LipoP1* in bundled and marginal patterns was significantly higher than in uncolonized regions. Data are means ± SD (n = 9 images from 6 plants). Different letters indicate statistically significant differences (p < 0.0001, ANOVA with Tukey test). (E) Expression patterns of different *CBP60g* lines (22-1, 22-4, 22-18 and 22-23) at 24hpi. Two-week old *Arabidopsis pCBP60g::NLS-3xmCitrine* seedlings were flood-inoculated with mCherry-tagged *Pst* DC3000. Pictures are maximum projections from confocal Z stacks of mCitrine, mCherry and chlorophyll autofluorescence signals. Chlorophyll autofluorescence is shown in gray. Scale bars: 20 µm

---

**Movie S1.** The immune marker *FRK1* is induced in surrounding cells of adaxial substomatal cavities colonized by *Pseudomonas syringae* DC3000. Two-week old *Arabidopsis pFRK1::NLS-3xmVENUS* seedlings were flood-inoculated with mCherry-tagged *Pst* DC3000. Movie of a Z stack showing brightfield, mVENUS and mCherry signals from the adaxial upper epidermis to mesophyll layer. **Related to** Figure 5A.

**Movie S2.** The immune marker *FRK1* is induced in surrounding cells of abaxial substomatal cavities colonized by *Pseudomonas syringae* DC3000. Two-week old *Arabidopsis pFRK1::NLS-3xmVENUS* seedlings were flood-inoculated with mCherry-tagged *Pst* DC3000. Movie of a Z stack showing brightfield, mVENUS and mCherry signals from the abaxial lower epidermis to mesophyll layer. **Related to** Figure 5A.

## SUPPLEMENTAL TABLE TITLES AND LEGENDS

**Table S1. - Differentially expressed genes in our studyRelated to Figure 1-2**.

**Table S2. Immunity and susceptibility genes used to calculate pathogen response score**. Related to Figure 2 **and STAR methods.**

**Table S3. Novel clusters of genes identified with pseudotime. Related to Figure 2 and S5.** Scores, including spatial autocorrelation score (Moran’s I), returned by Monocle3 for all genes in relation to Pseudotime. Genes with an associated module number (1-7) were identified as significantly correlated with Pseudotime.

**Table S4. Marker genes in each cluster. Related to STAR methods.** Per-gene marker statistics for all genes, determined by the FindAllMarkers function (Seurat). These describe the percentage of cells within (pct.1) and outside (pct.2) each cluster (seurat_clusters) that each gene is expressed, a p-value (p_val) for testing whether that gene is significantly enriched (Wilcox’s rank sum test), an adjusted p-value (p_val_adj; using the Benjamini-Hochberg method).

**Table S5. Primers used in this study. Related to STAR methods.**

**Table S6. Cluster names, types and cell metadata information. Related to STAR methods.** Assignments of cell clusters identified by Seurat to broad cell types/states. Per-cell metadata imputed for all cells in the integrated dataset. This includes transcript and gene counts for spliced, unspliced, and ambiguous mappings, number of transcripts derived from cholorplast (nCp) and mitochondrial (nMt) genes, the threshold UMI above which a cell was called from the raw counts matrix, the sample of origin (Sample_Name), statistics associated with SCT transformation, the predicted cell id and prediction scores for all cell types, a module score for pathogen responsiveness (DC3000.Up1 and DC3000.Down1), as well as cluster labels.

## Notes

### Competing Interest Statement

The authors have declared no competing interest.

