## Supplemental Table S2/5 for "Single-cell profiling of complex plant responses to *Pseudomonas syringae* infection"

**Table S2. Immunity and susceptibility genes used to calculate pathogen response score. Related to Figure 2.**

| Gene ID | Gene Symbol | Group |
|---|---|---|
| AT5G26920 | *CBP60g* | Immunity |
| AT4G04490 | *CRK36* | Immunity |
| AT4G23150 | *CRK7* | Immunity |
| AT4G39030 | *EDS5* | Immunity |
| AT2G19190 | *FRK1* | Immunity |
| AT4G11170 | *RMG1* | Immunity |
| AT5G52810 | *SARD4* | Immunity |
| AT3G11340 | *UGT76B1* | Immunity |
| AT2G14610 | *PR1* | Immunity |
| AT5G13320 | *PBS3* | Immunity |
| AT2G13790 | *SERK4* | Immunity |
| AT2G37710 | *LECRK-IV.1* | Immunity |
| AT2G42360 | *ATL41* | Immunity |
| AT2G39530 | *CASPL4D1* | Immunity |
| AT5G23820 | *ML3* | Susceptibility |
| AT3G46510 | *PUB13* | Susceptibility |
| AT1G19670 | *CORI1* | Susceptibility |
| AT4G23600 | *CORI3* | Susceptibility |
| AT5G24530 | *DMR6* | Susceptibility |
| AT3G28007 | *SWEET4* | Susceptibility |
| AT2G34070 | *TBL37* | Susceptibility |
| AT1G32640 | *JIN1/MYC2* | Susceptibility |
| AT4G12720 | *NUDT7* | Susceptibility |
| AT4G26080 | *ABI1* | Susceptibility |
| AT5G39670 | *CML46* | Susceptibility |

**Table S5. Primers used in this study. Related to Figures 3 and 4.**

| TAIR/AGI | Group | Gene symbol | Direction | Sequence (5'-3') | Source |
|---|---|---|---|---|---|
| AT3G18250 | Immune | *LipoP1* | Forward | CCCCTTCACCAGAAGCTGTACAAGAAAGTCG | This study |
| | | | Reverse | CTCTTCTTCTTTGGCATCTCTCTTTCTCTAGTTAATGTGG | |
| AT5G26920 | Immune | *CBP60g* | Forward | GCCCCCTTCACCTGGCTCGATCAAACTTAGATATCAATC | This study |
| | | | Reverse | CTTCTTCTTTGGCATTGATCACTTTTAGGTTTAGAG | |
| AT1G26770 | Susceptible | *EXPA10* | Forward | GCCGCCCCCTTCACCCGTATAGATAATAATTAAATGAATC | This study |
| | | | Reverse | CTCTTCTTCTTTGGCATGGGTGATATAAAATCAATTACTT | |
| AT4G00430 | Susceptible | *PIP1;4* | Forward | CCCCCTTCACCCGAACTTTACTAGTTACTATCTACC | This study |
| | | | Reverse | TTCTTCTTTGGCATTTCTCTCCCTCTCCCTC | |
| AT1G51780 | Susceptible | *ILL5* | Forward | CGCCCCCTTCACCTCTCACAGTATTAG | This study |
| | | | Reverse | CTTCTTCTTTGGCATCGTGAATCAAGAGATTGC | |
